# Immune profiling and treatment cessation provide mechanistic insights into disease pathogenesis and considerations for translating pre-clinical strategies in Leigh syndrome

**DOI:** 10.64898/2026.08.14.744649

**Authors:** Elizaveta A. Olkhova, Ernst-Bernhard Kayser, Anastasia Dimitriou, Michael Mulholland, Holly Coulson, Vivian Truong, Owen Cairns, Katerina James, Brittany M. Johnson, Monika Winter, Vandana Kalia, Sarkar Surojit, Allison Hanaford, Simon C. Johnson

## Abstract

Genetic mitochondrial diseases (GMDs) are major challenges to human health accounting for a significant fraction of heritable neurologic diseases, myopathies, and inborn errors of metabolism. Leigh syndrome (LS) is the most common clinical presentation of GMD in pediatric patients. LS is a severe and complex disease for which effective clinical therapies are currently lacking. Preclinical therapies identified in the *Ndufs4*(-/-) mouse model of LS include immune-targeting interventions and chronic mild hypoxia (11% oxygen). Immune-targeting interventions include rapamycin and high-dose pexidartinib, the latter appearing to fully suppress disease. The mechanisms underlying the benefits of hypoxia remain unclear, and the relationship between hypoxia and immune interventions have not been assessed. Here, we report the immune profile of brainstem of the *Ndufs4*(-/-) mouse model prior to and after disease onset and the impact of pexidartinib treatment. We provide evidence that macrophages/monocytes drive pathology, consistent with recent genetic studies. We additionally find that pre-disease onset animals lack signs of inflammation, and that the elimination of leukocytes fully suppresses the molecular signature of disease. Finally, using distinct post-developmental periods of treatment, we find pexidartinib and rapamycin provide benefits which persist long beyond treatment cessation, while cessation of hypoxia results in rapid disease onset and an acceleration of disease progression. These findings are consistent with hypoxia acting upstream of immune cell activation and have major implications for the therapeutic translation of both hypoxia and immune targeting interventions. Our findings establish hypoxia-cessation as a novel method for synchronizing inflammatory disease onset in the *Ndufs4*(-/-) model which will be useful in future mechanistic studies.

## Introduction

Genetic mitochondrial diseases (GMDs) are estimated to impact roughly 1 in 4,000 individuals and are extremely genetically and clinically heterogenous [1]. Subacute necrotizing encephalomyelopathy, or Leigh syndrome (LS), is the most common pediatric presentation of GMD. Patients with LS are often born without any overt signs of disease. Symptoms typically arise during the first years of life, though late-onset LS cases have also been reported [1].

LS is a severe and complex clinical entity. Patients typically experience neurologic and metabolic symptoms. While symptoms can vary from patient to patient, symmetric, progressive, bilateral neuroinflammatory lesions in specific brain regions, including the brainstem and basal ganglia, are a defining feature of the disease. These lesions can be detected via magnetic resonance imaging (MRI), facilitating diagnosis. Respiratory failure caused by the brainstem lesions is the proximal cause of death in most patients [2].

GMDs broadly, and LS specifically, are genetically heterogeneous: over 350 unique genes have been linked to GMD, with over 120 unique genes causally linked to LS alone [1]. These are encoded by both the mitochondrial (∼30% of patients) and nuclear (∼70% of patients) genomes, with both autosomal and X-linked genes. Nuclear gene origin GMDs can present as autosomal recessive, autosomal dominant, X-linked, or via compound heterozygosity. Disease penetrance and presentation in patients with pathogenic variants in mitochondrial DNA (mtDNA) is impacted by heteroplasmy levels both systemically and in key target tissues. Penetrance can be strongly impacted by genetic and environmental factors, often poorly understood.

Given this heterogeneity, any gene-specific interventions would be capable of benefitting only a small fraction of patients, limiting the populations that could benefit from approaches such as gene therapy (see ***Discussion***). Accordingly, identifying therapeutic strategies that can interrupt common pathways in disease pathogenesis downstream of multiple causal variants is considered critical for treating LS.

Our understanding of the pathogenesis of LS has advanced significantly in recent years, facilitated by the *Ndufs4*(-/-) transgenic mouse model of this disease [3]. This mouse carries a homozygous recessive loss of *Ndufs4* which encodes mitochondrial NADH dehydrogenase [ubiquinone] iron-sulphur protein 4 (Ndufs4), a structural/assembly component of mitochondrial electron transport chain complex I (ETC CI). The *Ndufs4*(-/-) mouse is a robust model of LS: loss of function variants in human homologue *NDUFS4* can cause LS in humans [1, 4–11], and the phenotype of *Ndufs4*(-/-) mice closely resembles human LS. Similarly to human patients, *Ndufs4*(-/-) mice are born without overt disease, first present with symptoms at ∼P37 (postnatal day 37), developing ataxia, metabolic dysfunction, seizures, progressive weight loss (cachexia), and showing a severely reduced lifespan. Critically, *Ndufs4*(-/-) mice develop bilateral progressive neuroinflammatory lesions in the brainstem consistent with those in humans [12–16]. As in humans, these drive respiratory dysfunction, reported to be the proximal cause of death if animals have not been euthanized for reaching defined endpoints [17].

Critically, studies using cell-type specific knockout of *Ndufs4* in neurons (Nestin-Cre) or neuron sub-populations have demonstrated that LS disease is driven by mitochondrial dysfunction in neuronal populations [12, 18, 19]. The entire LS phenotype of the global knockout is observed in the Nestin-Cre (pan-neuronal) driven knockout, while the majority of symptoms, including the progressive necrotizing brainstem lesions, are also present in a vGlut2-Cre (glutamatergic neuron specific) model [18, 19]. In the vGlut2-Cre driven knockout seizures are absent but severe fatal seizures occur in GABAergic neuron specific (Gad2-Cre driven) knockout [18, 19].

Through a series of recent studies, we’ve demonstrated that LS is an innate immune-mediated disease, that CNS lesions are driven by both microglia and peripheral macrophages/monocytes, and that the adaptive immune system plays no significant role in disease pathogenesis [13–15, 20]. Most strikingly, treatment with high dose pexidartinib, a colony-stimulating factor-1 receptor (Csf1r) inhibitor which depletes leukocytes, prevents LS symptoms in *Ndufs4*(-/-) mice [13]. Treated mice were generally free from LS-like symptoms, and the robustly (∼400%) increased survival was limited by drug toxicity of chronic pexidartinib treatment rather than disease. This finding represents the most robust pharmacologic intervention in the *Ndufs4*(-/-) model to date by a large margin. Other pharmacologic and genetic interventions targeting immune pathways also attenuate disease course including PI3Kγ inhibition, PKC inhibition, mTOR inhibition, and IFNγ deletion, though the benefits are modest compared to those of high-dose pexidartinib [13–15, 20, 21]. While these data demonstrate innate immune populations drive LS, many questions remain unanswered. The precise cell types, and relevant immune pathways, remain unresolved and may represent more precise therapeutic targets. In addition, while symptoms present postnatally in mice (∼P37) and humans (usually within the first years of life), whether subclinical neuroinflammation is present prior to symptom onset has not been assessed.

Similarly to high-dose pexidartinib, chronic mild hypoxia (11% O_2_) dramatically increases lifespan and suppresses CNS lesions and disease in the *Ndufs4*(-/-) model [22, 23]. Phlebotomy. carbon monoxide, and the small molecule HypoxyStat, which induces a hypoxia-like condition *in vivo* by altering the affinity of hemoglobin for oxygen, also attenuate disease in the *Ndufs4*(-/-) mouse model [24, 25]. The benefits of hypoxia are independent of hypoxia-inducible factor 1 alpha (Hif-1α) [24]. While chronic mild hypoxia is robust in the *Ndufs4*(-/-), the mechanisms underlying the benefits remain unclear, representing a knowledge gap that may impact efforts to translate this therapy into clinical use.

Given the robust effects of immune depletion and chronic hypoxia it is crucial to understand the relationship between these interventions. In addition, fundamental considerations regarding clinical translation remain unresolved. In particular, any risks associated with interruptions in or cessation of treatment have not been assessed.

Here, we assess the molecular signature immune activation in the *Ndufs4*(-/-) mouse brainstem tissue and examine disease onset and progression following treatment termination in the setting of mTOR inhibition, pexidartinib treatment, and hypoxia. We additionally establish hypoxia synchronization as a method for probing early mechanisms of immune activation and studying disease onset in the *Ndufs4*(-/-) model. These data provide new insights into the relationships between immune targeting and chronic hypoxia therapies and highlight critical considerations for clinical translation of these interventions.

## Results

### An extensive inflammatory signature develops in *Ndufs4*(-/-) brainstem after disease onset and is fully suppressed by pexidartinib

Neuroinflammatory lesions in the brainstem are a key defining feature of LS which drive many clinically important symptoms. To explore the inflammatory signature of *Ndufs4*(-/-) brainstem tissues and determine whether signs of inflammation are present prior to neurological symptom onset, we collected brainstem samples from control and *Ndufs4*(-/-) mice at postnatal days 23 (P23) and 45 (P45) for analysis via a NanoString mouse Pan Cancer Immune panel (see ***Fig. 1A, Methods***). These ages represent early post-weaning, pre-disease onset, and early post-disease, respectively. High-dose (300 mg/kg/d) pexidartinib prevents LS symptoms in the *Ndufs4*(-/-) model [13]. To assess the impact of this treatment on brainstem inflammation samples were also collected from pexidartinib treated mice at P45.

**Figure 1.**
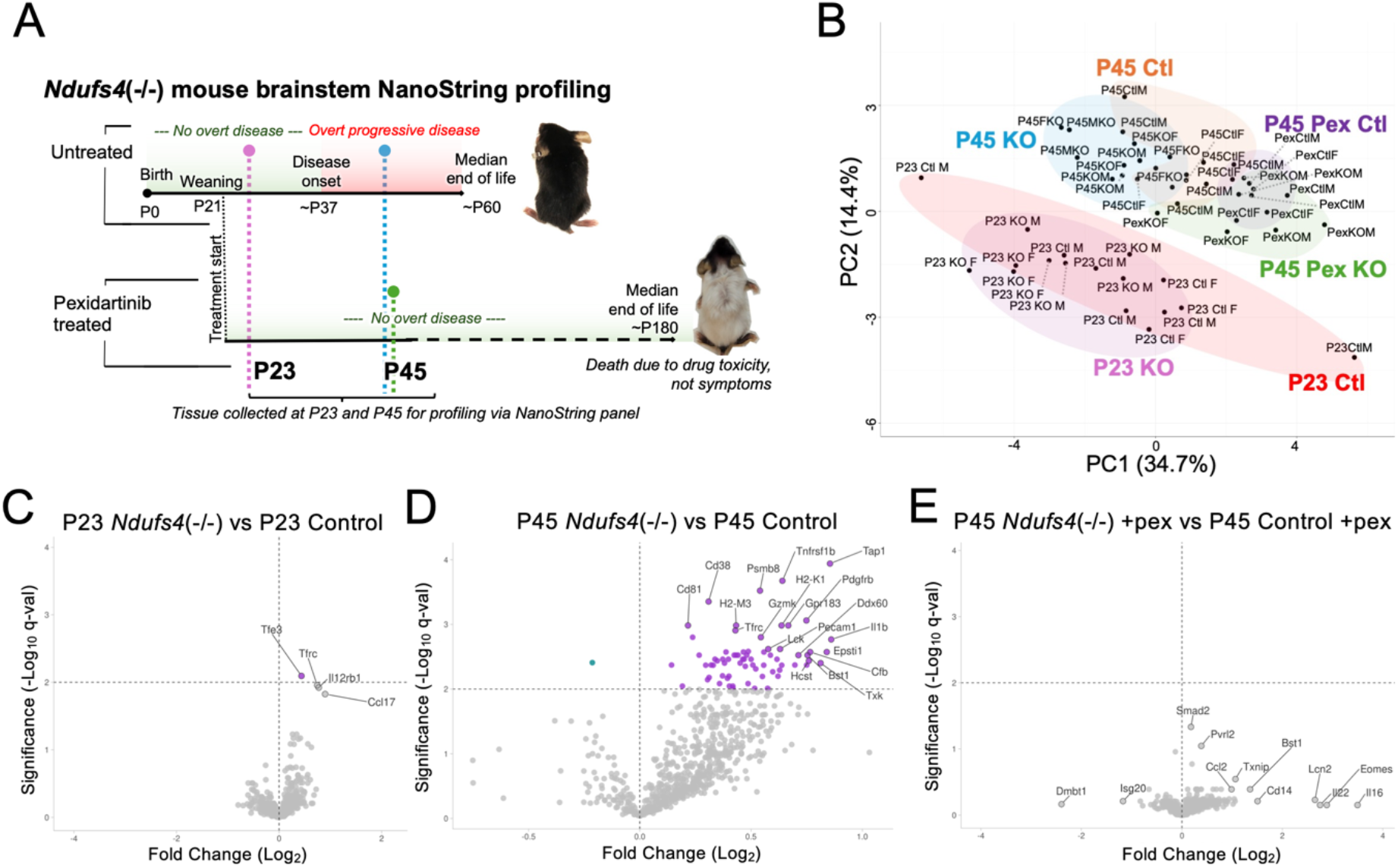
Extensive inflammatory changes develop after post-natal day 23 and are prevented by pexidartinib in the *Ndufs4*(-/-). (A) Overview of disease in *Ndufs4*(-/-) mice, treatment period for pexidartinib, and ages of tissue collection for NanoString analysis. Samples were collected from untreated (control diet) *Ndufs4*(-/-) and control mice at the post-weaning, pre-disease onset, age of post-natal day 23 (P23) and the overt but not end stage disease age of P45. Pexidartinib treated animal samples were collected at P45 (see ***Methods***). (B) Principal component (PC) analysis biplot of brainstem immune panel data (NanoString, see ***Methods***) from *Ndufs4*(-/-) and control mice at ages and treatments indicated in (A). At P23 control and *Ndufs4*(-/-) mouse cohorts show complete overlap in biplot distribution, suggesting a high degree of similarity prior to disease onset, compared to a greater degree of separation between P45 control and *Ndufs4*(-/-) animals. Pexidartinib treatment results in distinct clustering for both control and *Ndufs4*(-/-) cohorts, which partially overlap. (C-E) Volcano plots showing Log_2_(fold change) versus -Log_10_(q-value)(horizontal line at q=0.01). In each panel, the top right and left quadrants represent factors significantly elevated or reduced, respectively, in *Ndufs4*(-/-) compared to control animals in the given treatment. (C) Volcano plots of P23 control vs P23 *Ndufs4*(-/-) from animals in (B). Only one factor, *Tfe3*, is significantly increased in *Ndufs4*(-/-) mouse brainstem at P23 (see ***Results***). (D) Volcano plot of NanoString analysis of brainstem from P45 control vs P45 *Ndufs4*(-/-) from animals in (B). In contrast to P23, NanoString profiling reveals a widespread upregulation of inflammatory/immune factors in P45 *Ndufs4*(-/-) versus P45 control animals (see ***Results***, Fig. 2). (E) Volcano plot of NanoString analysis of brainstem from untreated control versus pexidartinib treated P45 *Ndufs4*(-/-) mice. No targets showing significant differences in expression between *Ndufs4*(-/-) and control animals (see ***Results***/***Discussion***).

Principal component analysis (PCA) of the NanoString dataset, which includes ∼750 immune/inflammatory transcripts, reveals a broad overlap between control and *Ndufs4*(-/-) samples at P23, consistent with the lack of observable phenotypes at this age (***Fig. 1B***). All P45 animals are separated from P23 animals in PCA clustering, suggesting significant shifts in immune/inflammatory gene expression as a function of age post-weaning. Untreated *Ndufs4*(-/-) and control mice form generally independent clusters at P45, while pexidartinib treated control and *Ndufs4*(-/-) mice largely overlap in their PCA distribution.

At the pre-disease onset (P23) only one panel transcript, *Tfe3*, is significantly upregulated in *Ndufs4*(-/-) compared to control mice by multiple-testing corrected q-value (FDR 1%, see ***Methods***)(***Fig. 1C***). In contrast, 70 of 750, nearly 10%, of the targets in the pan-immune panel, are significantly differentially expressed in the *Ndufs4*(-/-) versus control at P45 (***Fig. 1D***). Of these, all but one are increased in the *Ndufs4*(-/-).

Pexidartinib treatment fully suppressed the immune signature of disease at P45: no panel targets are significantly differentially expressed between pexidartinib treated control and *Ndufs4*(-/-) animals at P45 (***Fig. 1E***).

### Subclinical innate immune activity may occur as early as P23

Focusing first on inflammatory changes in the brainstem of the *Ndufs4*(-/-), we explored the differentially expressed genes in *Ndufs4*(-/-) versus control animals. As noted above, at P23 only one transcript is significantly upregulated in *Ndufs4*(-/-) mice compared to age matched controls – *Ttef3* (***Fig. 2A***). *Tfrc* and *Il12rb1*, and *Ccl17* additionally reach nominal (uncorrected) significance and border multiple testing corrected significance. *Ttef3* encodes a transcription factor highly expressed in neutrophils, while each of these factors are enriched among innate immune cell populations (see ***Fig. S1***)[26–28]. Together, these provide some evidence for subclinical innate immune activation prior to symptom onset (see ***Discussion***).

**Figure 2.**
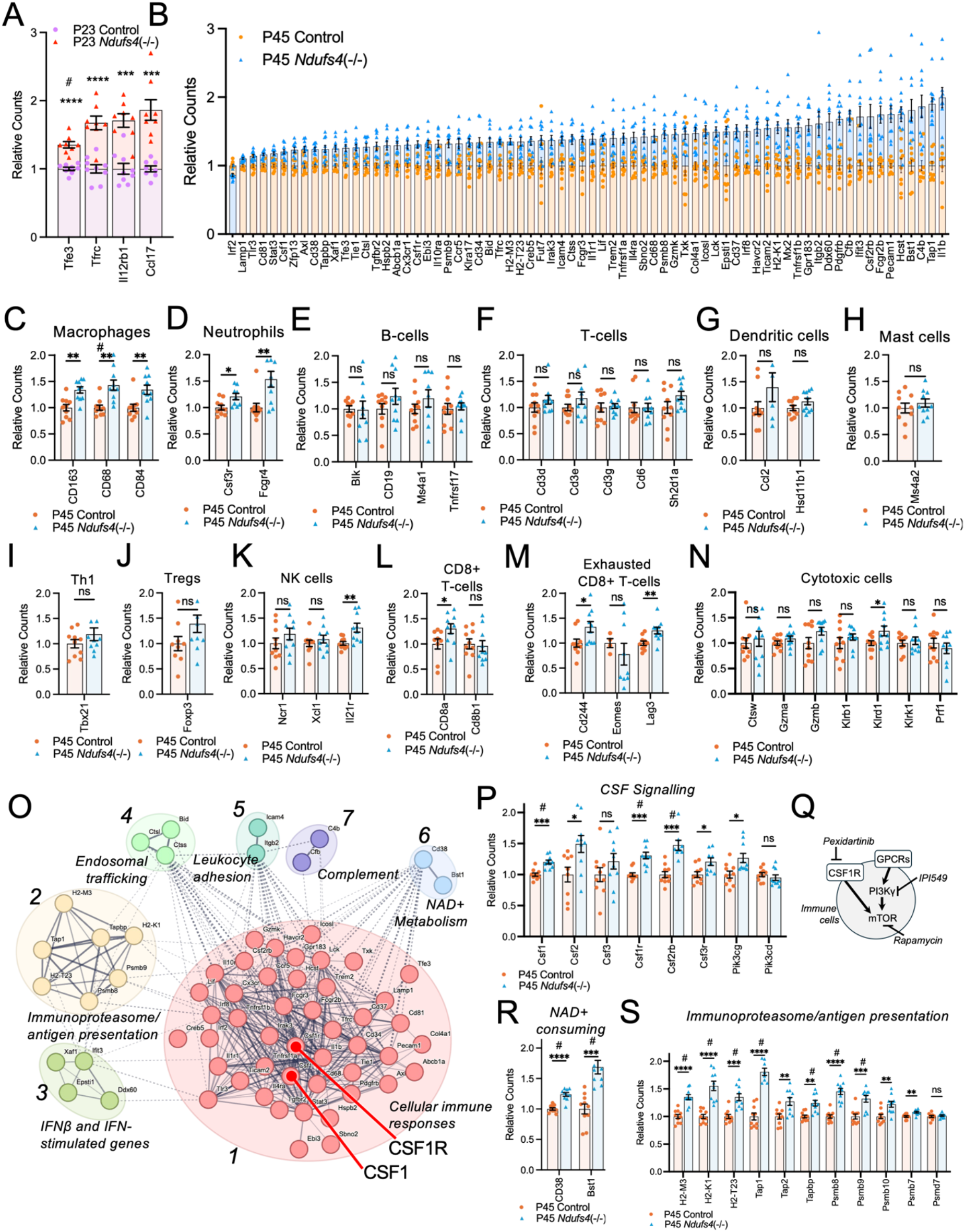
Immune profiling of *Ndufs4*(-/-) brainstem lysates reveals cellular mediators and disease-associated inflammatory processes. (A) Relative counts for immune panel targets upregulated at the pre-overt disease age of P23 (see volcano plot in Fig. 1C*, Methods*). Only one target, *Tfe3*, reached statistical significance following multiple testing correction (FDR<1%, see *Methods*), while three others were similarly upregulated and borderline significant (*Tfrc*, *Il12rb1*, *Ccl17*). All reached significance by uncorrected t-test. # - significant after multiple testing correction (FDR 1%, see *Methods*); ***p<0.0005, ****p<0.0001 by uncorrected pairwise t-test. (B) Relative counts for immune panel targets identified by FDR discovery to be downregulated (*Irf2*) or upregulated (all others plotted) in *Ndufs4*(-/-) versus control brainstem at the early post disease onset age of P45 (see volcano plot in Fig. 1D*, Methods*). (C-N) Relative counts in P45 *Ndufs4*(-/-) and control samples for NanoString panel markers associated with specific immune cell populations: (C) macrophages, (D) neutrophils, (E) B-cells, (F) T-cells, (G) dendritic cells, (H) mast cells, (I) T_h_1 helper T-cells, (J) Regulatory T-cells (Tregs), (K) natural killer (NK) cells, (L) CD8(+) T-cells, (M) exhausted CD8+ T-cells, and (N) cytotoxic cells (see *Methods* for NanoString cell type marker details). # - significant after multiple testing correction (FDR 1%, see *Methods*), **p<0.005, *p<0.05 by pairwise t-test. Error bars – SEM. (O) STRING (Search Tool for the Retrieval of Interacting Genes/Proteins) network clustering of targets with significantly increased expression in *Ndufs4*(-/-) brainstem at P45 versus age-matched controls (those transcripts in (B)). Protein-protein interaction (PPI) enrichment p-value = *0.038 (significant) versus background list (NanoString cancer pan-immune panel, see *Methods*). STRING cluster names: cluster 1 – cellular responses to molecules of bacterial origin; cluster 2 – antigen processing and presentation; cluster 3 – mixed including cellular responses to IFNβ and antiviral mechanisms by IFN-stimulated genes; cluster 4 – trafficking and processing of endosomal TLR, cathepsin propeptide inhibitor domain; cluster 5 – ICAM/ITGB2; cluster 6 – nicotinate and nicotinamide metabolism and NAD+ nucleotidase, cyclic ADP-ribose generating; cluster 7 – activation of C3 and C5, classical complement pathway C3/C5 convertase complex. (P) Select transcripts in the CSF signalling cluster that have previously been linked to the pathogenesis of LS via pharmacologic intervention studies in the *Ndufs4*(-/-) model (see text). #q<0.01 (multiple testing corrected), ***p<0.0005, *p<0.05 by pairwise t-test. Error bars – SEM. (Q) A simplified schematic representation of the relationship between previously established pharmacologic targets for attenuating disease in the *Ndufs4*(-/-) model which appear in (P). CSF1R (colony stimulating factor 1 receptor), PI3Kγ (phosphatidylinositol 3-kinase subunit gamma, encoded by *Pik3cg*), and mTOR (the mechanistic target of rapamycin). CSF1R, inhibited by pexidartinib, is necessary for leukocyte survival. PI3Kγ is specifically expressed in immune cells and mediates intracellular responses to extracellular signals, primarily downstream of GPCRs. mTOR is an intracellular signalling hub and a target for immune-modulating drugs such as rapamycin (see *Results* and *Discussion*). (R) Relative counts for NAD+ consuming ecto-enzymes (cluster 6 in O) upregulated in *Ndufs4*(-/-) versus control brainstem at P45. # - significant in discovery (FDR 1%). ****p<0.0001 by pairwise t-test. Error bars – SEM. (S) Relative counts for transcripts involved in the immunoproteasome and antigen presentation, including those in cluster 2 in (O) (*H2-M3*, *H2-K1*, *H2-T23*, *Tap1*, *Tapbp*, *Psmb8*, and *Psmb9*) as well as related transcripts (*Tap2*, *Psmb10*, *Psmb7*, *Psmd7*). #q<0.01 (multiple testing corrected); ****p<0.0001, ***p<0.0005, **p<0.005 by pairwise t-test. ns – not significant. Error bars – SEM.

### Broad immune panel upregulation is observed at the early post symptom onset age P45

*Irf2*, encoding a repressor of type-I interferon signaling is the only transcript significantly downregulated in *Ndufs4*(-/-) versus control at P45 [29], (***Fig. 2B***, far left). In contrast, 69 transcripts are significantly upregulated in the *Ndufs4*(-/-) (***Fig. 2B***).

### Innate, but not adaptive, immune cell markers are increased in *Ndufs4*(-/-) brainstem at P45

NanoString provides gene lists for transcripts enriched in individual immune cell types (see ***Supplemental file ‘****Mouse PanCancer Immune Panel’*). Among targets associated with macrophages and monocytes all are increased in *Ndufs4*(-/-) versus control brainstem at P45 (***Fig. 2C-D***). One, the monocyte lineage and tissue macrophage marker *Cd68*, reaches discovery threshold significance (FDR 1%), while all are significantly increased by unadjusted p-value.

In contrast, no B-cell, T-cell, dendritic cell, mast cell, Th1, or Treg markers are not increased in the *Ndufs4*(-/-)(***Fig. 2E-J***). Among NK, CD8+, exhausted T-cell, and cytotoxic cell markers a subset reach nominal significance (uncorrected p-value <0.05), all increased in the *Ndufs4*(-/-)(***Fig. 2K-N***). Overall, these data appear consistent with recent findings demonstrating that innate immune cells, including peripheral mononuclear phagocytic cells, drive CNS lesions while the adaptive immune system plays no role in disease in the *Ndufs4*(-/-)[13, 15](see ***Discussion***).

### Immune profiling identifies upregulated immune pathways

To identify any specific functional pathways involved in early post-disease onset pathology in LS we performed STRING (Search Tool for the Retrieval of Interacting Genes/Proteins) protein-protein interaction analysis of the 69 transcripts upregulated in *Ndufs4*(-/-) brainstem at P45 (see ***Methods***)[30]. Upregulated transcripts were significantly enriched for protein-protein interactions (PPI, *0.038 against the background NanoString Pan Cancer Immune panel list, see ***Methods***), and STRING analysis identified 7 clusters by Markov Cluster Algorithm (***Fig. 2O***). Key among these is a large central cluster composed of 46 transcripts and defined as *cellular responses to molecules of bacterial origin* (***Fig. 2O, cluster 1***).

*Csf1r*, which encode the target of pexidartinib (Csf1r), its agonist *Csf1*, and *Csf2rb* all appear in cluster 1 (see ***Discussion***). A broader examination of CSF1/2/3-related signaling factors beyond those which reached the discovery threshold (FDR 1%, see ***Methods***) reveals *Csf2*, *Csf3r*, and *Pik3cg* as elevated in the *Ndufs4*(-/-) brainstem at P45 by uncorrected p-value; in contrast, *Pik3d* is not elevated (***Fig. 2P***). These data appear consistent with the previously reported benefits of Csf1r, PI3Kγ, and mTOR inhibitors, and lack of benefits of PI3Kδ inhibition in the *Ndufs4*(-/-)(***Fig. 2Q***, see ***Discussion***)[13, 31–33].

Beyond this large central cluster six additional clusters included *antigen processing and presentation* (7 transcripts); *mixed including cellular responses to IFNβ and antiviral mechanisms by IFN-stimulated genes* (4 transcripts); and three additional smaller clusters involving endosomal TLR signaling, complement, and NAD+ metabolizing ectoenzymes. Two NAD+ degrading enzymes were identified as significantly increased in the *Ndufs4*(-/-) as P45 (***Fig. 2R***), a possible link between NAD+ metabolism and immune cell actions (see ***Discussion***).

Examining all immunoproteasome/antigen presentation transcripts including those reaching the discovery threshold (cluster 2) indicates the immunoproteasome, involved in presentation of intracellular antigens, is broadly upregulated in early post disease onset *Ndufs4*(-/-) brainstem (***Fig. 2S***). Non-standard histocompatibility complex subunits are also upregulated, including H2-M3, a subunit involved in presenting formylated peptides (bacterial or mitochondrial in origin, see ***Discussion***) [34].

### Pexidartinib effects place innate immune cells upstream of widespread inflammatory changes in the Ndufs4*(-/-)*

As detailed above, no genes in the NanoString immune panel were significantly upregulated in pexidartinib treated *Ndufs4*(-/-) compared to pexidartinib treated control mice at P45, a striking suppression of the widespread inflammation observed in untreated animals (see ***Fig. 1D***). These data suggest the majority of immune/inflammatory processes lie mechanistically downstream of pexidartinib depleted innate immune cells (see ***Discussion***).

To assess whether any immune/inflammatory pathways lie upstream of immune cell actions we assessed the impact of pexidartinib treatment within *Ndufs4*(-/-) animals, comparing pexidartinib and untreated *Ndufs4*(-/-) cohorts. Among the 750 gene set, 106 are significantly downregulated in pexidartinib treated versus control treated *Ndufs4*(-/-) at P45, while 7 are significantly upregulated (***Fig. 3A***). Transcripts upregulated in the pexidartinib cohort are candidates for processes not driven by immune cell activities, perhaps representing underlying CNS tissue or neuronal pathology (see ***Discussion***). Consistent with this notion, the top four include *Il1r1*, *Ccl11*, *Ppbp*, and *Vwf* (***Fig. 3A*, *Fig. 3B***), the latter three of which are most highly expressed in fibroblasts, platelets, and endothelial cells, rather than immune cell populations, in human scRNA datasets (***Figure S2, Discussion***).

**Figure 3.**
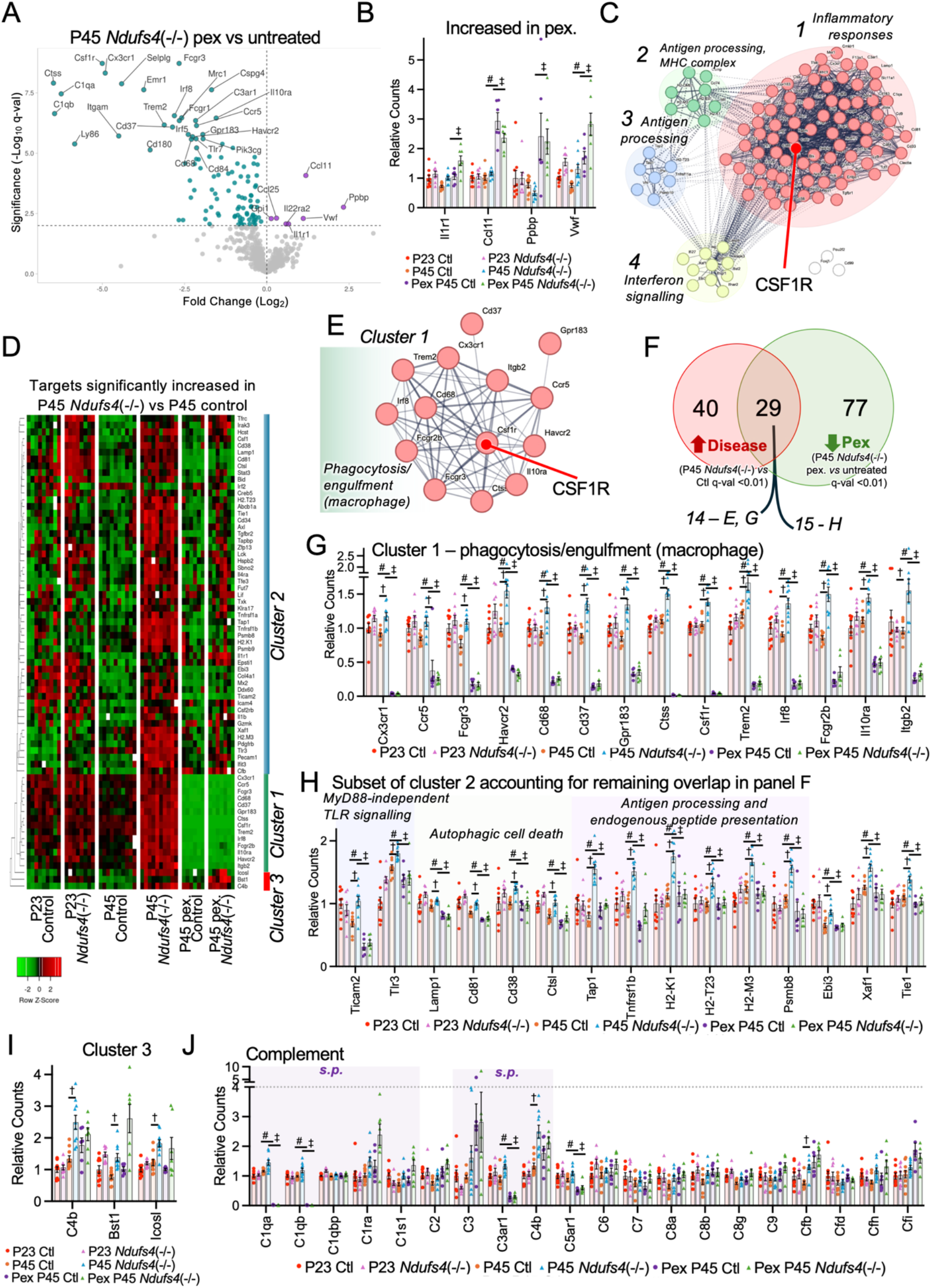
Immune panel targets downregulated by CSF1R inhibition suggest functional targets of treatment and putative upstream disease-initiating factors. (A) Volcano plots showing Log_2_(fold change) versus -Log_10_(q-value). Horizontal line at q=0.01. Top right and left quadrants represent factors significantly elevated or reduced, respectively, in pexidartinib treated *Ndufs4*(-/-) versus age-matched control treated *Ndufs4*(-/-) animals. (B) Scatter bar-plots of the four statistically significant most increased (by Log_2_(fold change) factors in pexidartinib treated *Ndufs4*(-/-) versus control treated *Ndufs4*(-/-) with all groups shown. # - P45 control versus P45 pexidartinib treated control significant by multiple testing corrected q-val (FDR 1%, see *Methods*). ‡ - P45 *Ndufs4*(-/-) versus P45 pexidartinib treated *Ndufs4*(-/-) significant by multiple testing corrected q-val (FDR 1%, see *Methods*). P23 data shown for reference. Uncorrected p-values not shown. Error bars – SEM. (C) STRING network clustering of targets with significantly lower expression in pexidartinib treated *Ndufs4*(-/-) versus control treated *Ndufs4*(-/-) at P45 (those transcripts in (A)). Protein-protein interaction enrichment ****p-value=1×10^-16^ versus the background set (Pan Cancer Immune profiling panel). STRING cluster names (first descriptor): cluster 1 – inflammatory response; cluster 2: interferon signalling; cluster 3: antigen processing and presentation of exogenous peptide antigens and (second descriptor) MHC protein complex; cluster 4 – antigen processing, cross presentation. (D) Hierarchical clustering (clustering performed using ClustVis, data displayed using Prism, see *Methods*) of the transcripts significantly increased in *Ndufs4*(-/-) versus control animals at P45 (targets in Fig. 2B) with all treatment groups included. (E) STRING analysis of cluster 1 in in the hierarchical heatmap from (D). Protein-protein interaction (PPI) enrichment p-value = ****3.17×10^-5^ versus background list (Pan Cancer Immune profiling panel, see *Methods*). (F) Overlap between transcripts significantly upregulated in *Ndufs4*(-/-) versus control at P45 (red circle, those in (D)) and transcripts significantly downregulated in pexidartinib treated versus control treated *Ndufs4*(-/-) at P45 (green circle). These 29 transcripts represent cluster 1 and a subset of cluster 2 in (D). (G) Relative counts for transcripts in cluster 1 (D, E). # - P45 control versus P45 pexidartinib treated control significant by multiple testing corrected q-val (FDR 1%, see *Methods*). ‡ - P45 *Ndufs4*(-/-) versus P45 pexidartinib treated *Ndufs4*(-/-) significant by multiple testing corrected q-val (FDR 1%, see *Methods*). † - P45 *Ndufs4*(-/-) versus P45 controls significant by multiple testing corrected q-val (FDR 1%, see *Methods*). P23 data shown for reference. Uncorrected p-values not shown. Error bars – SEM. (H) Relative counts for those transcripts in cluster 2 (D, E) that are also significantly lower in pexidartinib compared to control treated *Ndufs4*(-/-) animals at P45. # - P45 control versus P45 pexidartinib treated control significant by multiple testing corrected q-val (FDR 1%, see *Methods*). ‡ - P45 *Ndufs4*(-/-) versus P45 pexidartinib treated *Ndufs4*(-/-) significant by multiple testing corrected q-val (FDR 1%, see *Methods*). † - P45 *Ndufs4*(-/-) versus P45 controls significant by multiple testing corrected q-val (FDR 1%, see *Methods*). P23 data shown for reference. Uncorrected p-values not shown. Error bars – SEM. (I) Relative counts for cluster 3 (from panel D). † - P45 *Ndufs4*(-/-) versus P45 controls significant by multiple testing corrected q-val (FDR 1%, see *Methods*). P23 data shown for reference. Uncorrected p-values not shown. Error bars – SEM. (J) Relative counts for complement factor transcripts. # - P45 control versus P45 pexidartinib treated control significant by multiple testing corrected q-val (FDR 1%, see *Methods*). ‡ - P45 *Ndufs4*(-/-) versus P45 pexidartinib treated *Ndufs4*(-/-) significant by multiple testing corrected q-val (FDR 1%, see *Methods*). † - P45 *Ndufs4*(-/-) versus P45 controls significant by multiple testing corrected q-val (FDR 1%, see *Methods*). P23 data shown for reference. Uncorrected p-values not shown. Error bars – SEM.

### Profiling reveals pathways targeted by pexidartinib and potential disease initiating processes

To identify targets of pexidartinib treatment, we next assessed the transcripts significantly downregulated by pexidartinib. STRING profiling of downregulated transcripts identifies a significant enrichment of interactions (PPI enrichment ****p-value <1×10^-16^ against the background list) and three clusters (***Fig. 3C***). These include a large ‘inflammatory responses’ cluster similar to the cellular immune responses cluster in *Ndufs4*(-/-) versus control mic (***Fig. 2O***); antigen processing and antigen processing/MHC complex, overlapping with immunoproteasome and antigen processing in *Ndufs4*(-/-) versus control mice (***Fig. 2O***); and interferon signaling. As with the central cluster in control vs *Ndufs4*(-/-) mice (***Fig. 2O***), *Csf1r* is highly connected in the major cluster for pexidartinib-downregulated genes (***Fig. 3C***).

To gain further insights into the targets of pexidartinib which may be relevant to pathology in the *Ndufs4*(-/-) we focused on transcripts significantly increased in untreated P45 *Ndufs4*(-/-) mice (those in ***Fig. 2B***), assessing expression across all cohorts (***Fig. 3D***). Using unbiased clustering, transcripts from this group fall into three clusters based on patterns that can be broadly described as 1) upregulated in P45 *Ndufs4*(-/-) and completely ablated by pexidartinib treatment (Cluster 1); 2) upregulated in P45 *Ndufs4*(-/-) and returned to control levels by pexidartinib treatment (Cluster 2); and 3) a small set upregulated in P45 *Ndufs4*(-/-) but not changed by pexidartinib treatment (***Fig. 3D***).

Cluster 1 – genes upregulated in P45 *Ndufs4*(-/-) and ablated by pexidartinib – is composed of 14 transcripts. Encoded proteins comprise one large functional cluster by STRING analysis: phagocytosis/engulfment (macrophages), centered on Csf1r (***Fig. 3E***). These represent approximately half (14 of 29) of all transcripts overlapping between those upregulated in disease (P45 *Ndufs4*(-/-) versus control) and those downregulated by pexidartinib (P45 *Ndufs4*(-/-) pexidartinib treated versus untreated) (***Fig. 3F***). Each of these is potently suppressed by pexidartinib in both *Ndufs4*(-/-) and control animals, indicating they represent transcripts expressed by cell types eliminated by pexidartinib (***Fig. 3G***, see ***Discussion***).

Cluster 2 includes the remaining 15 targets in the pexidartinib/disease overlap (see ***Fig. 3F***). STRING analysis of this cluster identifies three functional interaction networks among this group – *MyD88*-independent TLR signaling, autophagic cell death, and antigen processing and endogenous peptide presentation via the immunoproteasome/MHC (***Fig. 3H***, see ***Discussion***).

Cluster 3, which includes transcripts upregulated in the P45 *Ndufs4*(-/-) and attenuated by pexidartinib, includes complement factor C4b, the NAD+ ectoenzyme Bst1, and the IFNγ-induced T-cell regulator *Icosl* (***Fig. 3GI***, see ***Discussion***).

Complement pathways are involved in cell-mediated inflammation and linked to the process of synaptic pruning (see ***Discussion***). Given the complement factors in cluster 3, we plotted the full set of complement factors represented in the NanoString Cancer Pan Immune panel (***Fig. 3J***). Of these, four follow the pattern of upregulation in the *Ndufs4*(-/-) at P45 (compared to control mice) and robust suppression by pexidartinib (C1qa, C1qb, C3ar1, and C5ar1). 10 are upregulated in *Ndufs4*(-/-) compared to control at P45 but remain elevated or show a further increase in elevation with pexidartinib treatment, consistent with a process mechanistically upstream of immune cell mediated disease. At least 4 of these 10 are involved in synaptic pruning (C1ra, C1s1, C3, C4b, in groups labeled *s.p.* in ***Fig. 3J***, see ***Discussion***).

### Disease progression following cessation of therapeutic interventions is consistent with hypoxia acting upstream of immune activation

The precise mechanisms mediating the beneficial effects of therapeutic chronic mild hypoxia in the *Ndufs4*(-/-) are unclear, though it is thought to act upstream of immune cell activation, in contrast with pexidartinib, which acts by depleting innate immune populations.

Our prior observations and the above NanoString data led to two specific hypotheses: 1) if hypoxia acts upstream of immune cell activation, cessation of hypoxia at a post-disease onset age should result in rapid immune activation triggering an abrupt disease onset as immune cells are present in circulation and can be quickly recruited to respond to any stress signal. 2) disease onset after immune depletion should be delayed while immune cell repopulation occurs.

To test these hypotheses, we treated *Ndufs4*(-/-) mice with chronic mild hypoxia (11% O_2_), 300 mg/kg/day pexidartinib, or 8 mg/kg/day rapamycin from P26-P40, spanning the disease onset age of∼P37 (see ***Methods***), and assessed disease outcomes. Following the cessation of treatment, mice that had been treated with either rapamycin or pexidartinib continue to gain weight for ∼10-14 days, followed by the onset of weight loss in pexidartinib treated mice and a long-term maintenance of weight in rapamycin treated mice (***Fig. 4A-B***). In contrast, animals treated with hypoxia start to lose weight within a few days of treatment cessation and their weight loss followed the trajectory of untreated animals (***Fig. 4A-B***). All but one hypoxia cessation animal reached their maximum weight by P43, three days after hypoxia cessation, with the last reaching a weight peak at P46.

**Figure 4.**
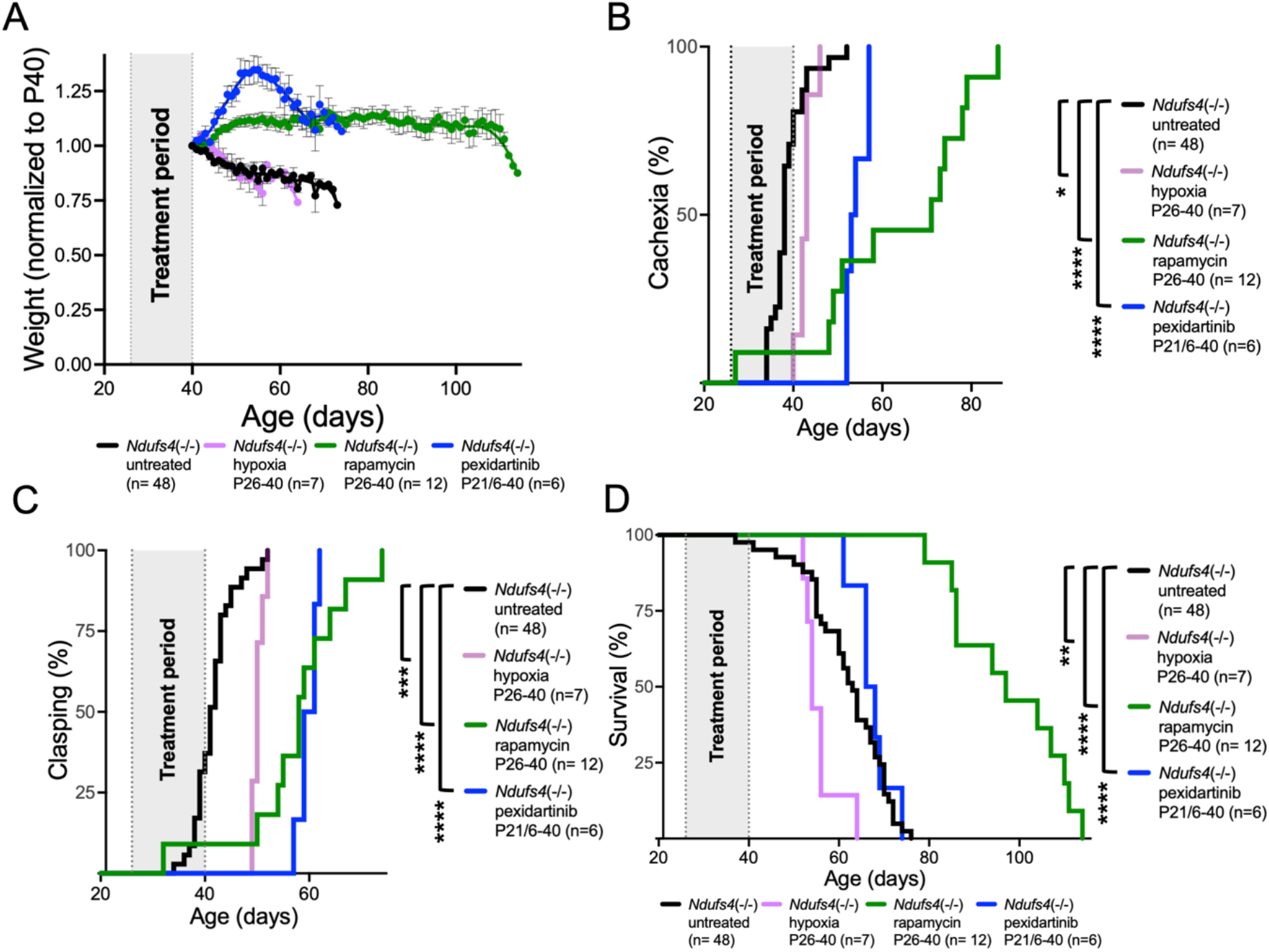
Benefits of immune-targeting interventions persist after treatment cessation while hypoxia cessation leads to rapid disease onset. (A) Weight normalized to P40 in *Ndufs4*(-/-) animals treated with rapamycin, pexidartinib, chronic mild hypoxia (8 mg/kg/day by IP injection, 300 mg/kg/day oral treatment, and 11% oxygen, respectively, see *Methods* and text for details) until P40 or provided no intervention (untreated). Error bars – SEM, datapoints are mean values, trend lines are LOWESS curves (see *Methods*). (B) Onset of cachexia (weight loss) in animals from (A). Data plotted represent percentage (%) of animals which have shown cachexia in each cohort by age. (C) Onset of clasping (early visually assessed neurologic disease sign, see *Methods*) in animals from (A). Data plotted represent percentage (%) of animals which have shown clasping in each cohort by age. (D) Survival of animals from (A). Data plotted represent percentage (%) of animals surviving by age. (A-D) *p<0.05, **p<0.005, ***p<0.0005, ****p<0.0001 by log-rank test, comparison pairs as indicated. Only relevant comparisons shown.

Clasping, a widely reported early sign of neurodegenerative disease onset in the *Ndufs4*(-/-) model, is similarly delayed in pexidartinib cessation and rapamycin cessation cohorts compared to hypoxia cessation, with median onset at 60, 58, and 50, respectively (***Fig. 4C***).

Brief treatment with rapamycin during an early developmental window provides a long-term benefit to survival - median survival in mice treated with rapamycin from P26-40 was 97 days versus 63 in untreated *Ndufs4*(-/-) animals (***Fig. 4D***). Median survival in P26-40 pexidartinib treated mice was slightly, but not significantly, longer than untreated animals (median P67 days). In contrast to both rapamycin and pexidartinib, hypoxia treatment from P26-40 was associated with a significant *reduction* in ultimate survival (median 54 days) (see ***Discussion***).

### Hypoxia cessation leads to a significant acceleration of disease course in the *Ndufs4*(-/-) mouse model of disease

To assess whether adjusting the treatment period beyond the immediate period of disease onset might alter the effects of hypoxia cessation we additionally treated mice with mild hypoxia from P21-40 and P21-60, following outcomes as above. In all hypoxia paradigms animals are healthy during hypoxia treatment, with P21-60 reproducing the benefits reported for this intervention up until treatment cessation (***Fig. 5A***). However, cessation of hypoxia led to rapid onset of weight loss and symptoms (***Fig. 5***). Notably, the rate of weight loss taken from the age of hypoxia removal was significantly increased in those animals treated for longer periods, and rate of weight loss was greater in all hypoxia cessation groups compared to untreated *Ndufs4*(-/-) mice (***Fig. 5A-C***).

**Figure 5.**
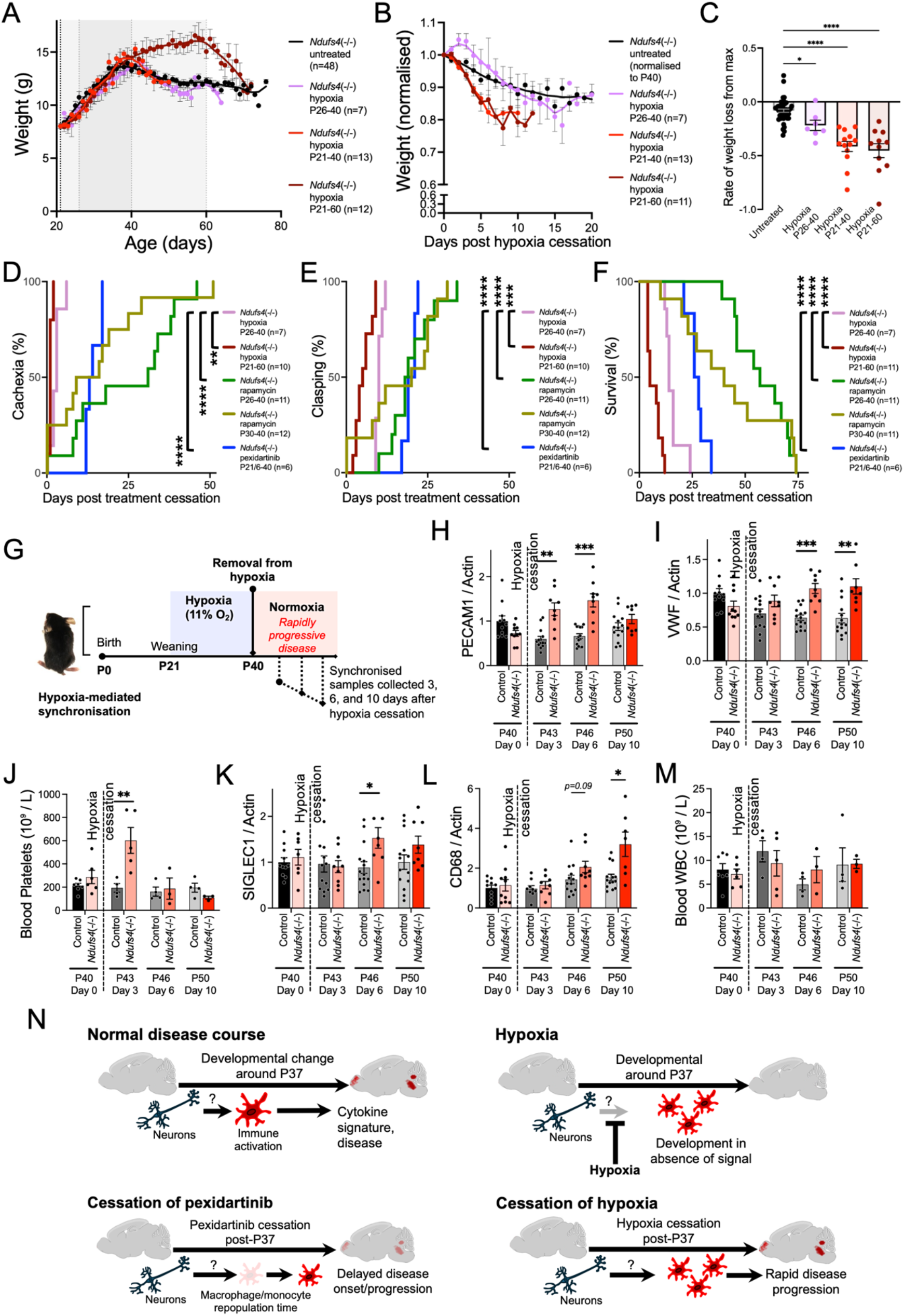
Hypoxia cessation leads to accelerated disease but provides an experimental strategy for disease onset synchronization. (A) Weight over time of *Ndufs4*(-/-) animals being treated with chronic mild hypoxia (11% oxygen, see *Methods* and text for details) from P26-40, P21-40, or P21-60, or housed in normoxia (untreated). Error bars – SEM, datapoints are mean values, trend lines are LOWESS curves (see *Methods*). (B) Plots of animal weights from panel (A) normalized to the day of hypoxia cessation. Error bars – SEM, datapoints are averages, trend lines are LOWESS curves (see *Methods*). (C) Rate of weight loss from the date maximum weight was recorded. Error bars – SEM, regular one-way ANOVA p<0.0001, *p<0.05, ****p<0.0001 by Dunnet’s multiple comparisons test. (D) Onset of cachexia (weight loss) in animals from (A). Data plotted represent percentage (%) of animals which have shown cachexia in each cohort by age. (E) Onset of clasping (early visually assessed neurologic disease sign, see *Methods*) in animals from (A). Data plotted represent percentage (%) of animals which have shown clasping in each cohort by age. (F) Survival of animals from (A). Data plotted are percent (%) animals surviving by age. (A-D) *p<0.05, **p<0.005, ***p<0.0005, ****p<0.0001 by log-rank test, comparison pairs as indicated, only relevant pairwise comparisons shown. None of the rapamycin P26-40 vs rapamycin P30-40 comparisons were significant. (G) Schematic of hypoxia removal as a method for synchronization of disease onset. (H-I) Expression, by qPCR analysis, of vascular markers *Pecam1* and *Vwf* from brainstem lysates of hypoxia synchronized animals (see *Methods*). (J) Change in blood platelets over time in hypoxia-synchronized animals. (K-L) Expression, by qPCR analysis, of macrophage-specific markers *Siglec1* and *Cd68* from brainstem lysates of hypoxia synchronized animals (see *Methods*). (M) Blood WBC (white blood cell) counts over time in hypoxia-synchronized animals. (N) Conceptual models for disease onset, the actions of hypoxia, and disease onset following pexidartinib or hypoxia cessation in the *Ndufs4*(-/-) mouse model (see *Discussion*).

The onset of symptoms, when considered as a function of time after treatment cessation, is significantly accelerated in *Ndufs4*(-/-) animals exposed to hypoxia compared to those treated with rapamycin or pexidartinib. Cachexia onset is apparent by the second day post cessation in *Ndufs4*(-/-) mice treated in hypoxia from P21-60, similar to the rapid onset noted above for treatment from P26-40 (***Fig. 5D***). In contrast, median cachexia onset was 12.5 days post cessation for pexidartinib treated *Ndufs4*(-/-) mice and 13.5 and 31 days post cessation for those treated with rapamycin from P26-40 (***Fig. 5D***). Cachexia onset was also delayed compared to hypoxia cessation animals in a cohort of *Ndufs4*(-/-) mice treated with rapamycin for an even shorter period - P30-40 (***Fig. 5D***).

Similarly, clasping onset was more rapid in hypoxia cessation mice compared to pexidartinib or rapamycin cohorts (***Fig. 5E***). Mice treated with hypoxia from P21-60 show the most rapid onset of clasping, with a median onset of 5 days post treatment cessation compared to 10 days for hypoxia treatment from P26-40, 20 days for pexidartinib P26-40 treated mice, and 18.5 and 20 for mice treated with rapamycin from P26-40 or P30-40, respectively.

Survival after treatment cessation was shortest in the hypoxia cohorts. As noted above, hypoxia treatment from P26-40 results in an *Ndufs4*(-/-) lifespan shorter than in untreated mice, with a median survival of 14 days post treatment cessation (***Fig. 5F***). This is in contrast with a median post-cessation survival of 27 days for mice treated with pexidartinib from P26-40. Hypoxia treatment from P21-60 is associated with even shorter post-cessation survival, with a median survival of only 5 days post-cessation (***Fig. 5F***). Treatment with rapamycin from P30-40 or P26-40 resulted in median post-cessation survival of 40 and 57 days, respectively. Given that the ultimate survival in animals treated with rapamycin from P26-40 was near that of animals treated with rapamycin for life [32] we did not add additional, longer duration, cohorts.

### Hypoxia cessation provides a method for synchronizing immune-mediated disease onset in the Ndufs4(-/-)

Given these findings, hypoxia-cessation appears to provide a useful experimental paradigm for synchronizing disease onset in the *Ndufs4*(-/-), enabling the identification of early and late factors involved in disease onset. As a proof of concept, we treated a large cohort of animals with hypoxia until P40, with animals either used for tissue collection at that age (directly from hypoxia) or moved to normoxic conditions and harvested at 3, 6, and 10 days post-cessation (***Fig. 5G***). We assessed the expression in brainstem of targets identified by NanoString (above) to be increased in P45 *Ndufs4*(-/-) and not attenuated by pexidartinib - PECAM1 and VWF - and or increased P45 *Ndufs4*(-/-) and rescued by pexidartinib - SIGLEC1 and CD68 (see above). Based on our data here and elsewhere these represent putative early (upstream of immune cell involvement) and later (immune cell markers) markers of disease pathogenesis (see ***Discussion***).

PECAM1 and VWF, expressed by platelets and endothelial cells, are both significantly increased in post-removal *Ndufs4*(-/-) versus control (at 3 and 6 days and 6 and 10 days, respectively), with *Ndufs4*(-/-) mice diverging from controls by 3 days post hypoxia (***Fig. 5H-I***). Intriguingly, *Ndufs4*(-/-), but not control, animals also show a spike in platelet numbers by complete blood cell (CBC) analysis (***Fig. 5J***, complete CBC data in ***Table S1,*** see ***Discussion***).

SIGLEC1 and CD68, markers of macrophages/monocytes, are also show significant increases in the post hypoxia cessation period, but the pattern of expression is delayed compared to PECAM1 and VWF (***Fig. 5L-M***). Notably, while these markers are elevated in brainstem lysates in the hypoxia-synchronized *Ndufs4*(-/-) mice compared to controls, there is no detectable change in white blood cell counts by CBC (***Fig. 5M, Table S1***).

## Discussion

Here, using NanoString profiling of immune-factors and *in vivo* treatment-cessation experiments, we provide novel insights into the pathogenesis of Leigh syndrome. We identify new therapeutic candidates for intervention, further our mechanistic understanding of existing pre-clinical therapeutic interventions, and reveal important considerations for the use of hypoxia-related treatments.

### Postnatal disease onset but some evidence for subclinical stress by P23

In human LS patients there is typically a disease-free postnatal window. The majority of LS diagnoses are in pediatric patients, with symptoms typically first appearing before the age of 2, though many late onset and adult onset LS and Leigh-like encephalopathy cases have also been reported [1, 35–38]. Early stages of disease onset are exceptionally difficult to study in humans due to the postnatal presentation of symptoms, delays in diagnosis, and the extreme genetic and clinical heterogeneity of the disease. Some evidence suggests that immune stressors can drive disease onset in humans (reviewed in [39]), but the mechanisms involved in disease onset remain largely unknown.

In the *Ndufs4*(-/-) mouse model of LS disease onset occurs postnatally at ∼P37. Prior to this age, animals tend to be slightly small for developmental age and present with a transient hair cycle phenotype but are otherwise free from an overt phenotype, including signs of disease [3, 12, 13, 32, 40]. While we previously demonstrated that immune cells depletion prevents symptoms [13], the landscape of CNS inflammation had not been reported, including at ages preceding observable symptoms or detectable CNS lesions. Here, we find here that there is no broad neuroinflammatory signal at P23, consistent with the lack of phenotype and broadly supportive of the notion that disease pathogenesis initiates after weaning. However, a few markers were increased at P23 (see ***Figures 1, 2***) suggest subclinical immune-related stress may be present even at this early age. Notably, these appear to implicate neutrophils as potential early participants in disease (see above). Neutrophils have recently been implicated as early drivers of a variety of CNS diseases, from neuromyelitis optica to Alzheimer’s, are acutely responsive to mitochondria-derived innate immune activating signals such as formylated peptides (see below), and may represent a viable early/upstream target in the pathogenesis of LS [35, 41–45].

### Innate immune cells drive the neuroinflammatory signature of LS

One key finding in this study is that CSF1R inhibition leads to a full suppression of the inflammatory signature of LS brainstem. This reveals that the bulk of inflammatory cytokines detected in LS are produced secondary to immune cell actions, rather than lying upstream of immune cell involvement. An alternative possibility was that depletion of CSF1R inhibitor sensitive immune cells would result in a residual signature of inflammation arising from other cell types. Given that it has been previously shown that neurons drive disease in LS (see ***Introduction***) this possibility seemed likely [18, 19].

While the cancer pan-immune panel is not all-inclusive, the 770 gene panel represents diverse immune cells and pathways. Accordingly, our data provide strong evidence that CSF1R responsive cells (mononuclear phagocytes, T-cells, B-cells, *etc.*) drive the neuroinflammatory signature of LS.

In addition, analysis of cell type markers indicates that innate immune cells – macrophages, monocytes, and neutrophils – are increased in the *Ndufs4*(-/-) brainstem at early post-disease onset age of P45, while B-, T-, and NK-cells are not. This is consistent with our recent prior body of work demonstrating that peripheral monocytes/macrophages drive LS pathology and that depletion of depletion of adaptive immune cells has no impact on disease [13–16].

Beyond providing additional supportive evidence that LS is an innate immune mediated disease, the NanoString datasets here implicate neutrophils. As noted above, neutrophils may act early in disease onset and might provide new targets for targeted interventions - perhaps providing disease attenuation with reduced off-target effects and more limited immune suppression.

### Differential effects of immune-targeting therapeutics

The identification of Csf1r as a central upregulated factor among the immune panel gene set in P45 *Ndufs4*(-/-) versus control brainstem lysates is consistent with our prior findings with CSF1R/PI3Kγ/mTOR inhibitors, directly linking our NanoString profiling data here to experimental outcomes in mice [13, 32, 33, 46].

Mechanistic explanations for the differential efficacies of distinct immune-targeting interventions have been lacking. Our data here provide additional context for interpreting prior immune intervention studies in mice and in human MD patients, for example the lack of efficacy of calcineurin inhibitors such as tacrolimus: we’ve shown that tacrolimus does not provide any benefits in the *Ndufs4*(-/-) mouse model, and in a small clinical study found mTOR inhibitors provide benefits to post-organ transplant MELAS (Mitochondrial Encephalopathy, Lactic Acidosis, and Stroke-like episodes) patients previously managed with tacrolimus or cyclosporin [32, 46]. A re-assessment of these findings in light of the cell type enrichment data presented here, as well as our recent work genetically targeting the adaptive immune system [15], suggests the efficacy of immune-targeting drugs is dependent on their relative impact on innate immune cell populations. Tacrolimus and cyclosporin act primarily through actions on T-cells, while mTOR and CSF1R inhibitors also impact innate immune functions [47, 48]. These findings will help identify and prioritize future candidates.

The profiling data here also provides a new hypothesis for the specific benefits of the PI3Kγ inhibitor IPI-549 [13]. PI3Kγ is an PI3K catalytic subunit isoform specifically expressed by myeloid cells, involved in neutrophil and macrophage activation and migration and driving classic M1 polarization and inflammation in some disease settings [49–51]. We’ve found the PI3Kγ inhibitor IPI-549 significantly attenuates disease in the *Ndufs4*(-/-), while inhibitors of other PI3K catalytic subunit isoforms do not [13]. Notably, PI3Kγ mediates signaling through GPCRs, including the FPRs (discussed below), consistent with a potential role for FPRs in disease pathogenesis [52, 53]. Accordingly, the unique benefits of IPI-549 versus other PI3K isoform inhibitors may be linked to its role in mediating FPR signaling.

Overall, our findings lead to the prediction that drugs primarily targeting adaptive immune populations will have limited benefits compared to drugs targeting innate immune populations, and that it may be possible to target the specific pathway(s) of induction. This represents an important step in developing relevant therapeutic approaches.

### NAD+ ectoenzymes

An intriguing finding was the significant upregulation of transcripts encoding NAD+ ectoenzymes Bst1 and CD38 among those significantly upregulated in the P45 *Ndufs4*(-/-) compared to control. CD38 is a transmembrane glycoprotein found on many immune cell types which acts as a primary consumer of NAD+ and regulator of NAD+ levels [54]. CD38-mediated depletion of NAD+ has been implicated in inflammation broadly, including in aging [55–57]. Bst1 is similarly highly expressed in immune cells and is thought to impact NAD+, though its role is less clear [58, 59]. NAD+ depletion is part of the underlying molecular defect in ETC CI (NADH dehydrogenase) defective mutants, such as the *Ndufs4*(-/-) model, and NAD+ precursor supplementation has been shown to modestly impacting disease outcomes in this model [60]. Accordingly, there are clear potential mechanistic links between these enzymes and disease pathology in the *Ndufs4*(-/-). First, they may exacerbate the primary molecular defect and downstream consequences (e.g. the dysregulation of NADH/NAD+ regulated pathways), driving disease. Second, their upregulation may be mechanistically responsible for the observed benefits of NAD+ precursor therapies, which modestly attenuated symptoms but did not fundamentally alter disease course. Further study of the role of these NAD+ ectoenzymes in the pathobiology of LS appears warranted.

### Candidates for disease-initiating pathways

While disease in the *Ndufs4*(-/-) is known to arise from neurons and be mechanistically driven by immune cells (see above), the neuron-derived signal driving immune activation (direct or indirect) is not yet known. Identifying the immune activating signal or signals is viewed as critical, as they may provide potent and specific therapeutic targets. The NanoString data implicate two candidates – formylated peptides and complement-mediated synaptic pruning.

N-formylated peptides, present in bacteria and mitochondria-derived proteins, but not nuclear-encoded proteins, are potent immune activators. Here we find genes involved in presentation of intracellular bacterial peptides are upregulated early in disease in *Ndufs4*(-/-) brainstem, including formylated-peptide induced non-classical major histocompatibility complex (MHC) class 1b molecule H2-M3, which uniquely specializes in presenting formylated peptides, and TAP1/TAP2, known to present formylated peptides derived from mitochondria-encoded ETC CI subunit ND1 [61–64].

FPR1 mediates the majority of pro-inflammatory signaling from formylated peptides, with downstream signaling mediated by PI3Kγ (see above), PKC (inhibitors of which also attenuate disease in the *Ndufs4*(-/-)), and other pathways [21, 62]. Inhibition of FPR1 has been shown to attenuate multiple diseases driven by innate immune cell infiltration of the CNS, making it a strong candidate for therapeutic intervention [65, 66].

We also find evidence for increased expression of transcripts associated with complement-mediated synaptic pruning. Aberrant synaptic pruning is linked to a variety of neurodevelopmental and neuroinflammatory diseases such as autism, Alzheimer’s, and multiple sclerosis [67–70]. Pruning is an important developmental process, providing an intriguing possibility that aberrant pruning may underlie the post-natal onset of disease in LS.

It is possible neither of these processes mechanistically drive LS, that they are both major contributors, or that they are two of multiple processes involved in disease. The role of FPR signalling and pruning must be assessed experimentally, but our data indicate they are strong candidates for further study.

### Treatment cessation places hypoxia upstream of immune activating signals and implications for therapeutic intervention

Chronic mild hypoxia and immune depletion via CSF1R inhibition with pexidartinib provide similar benefits in the *Ndufs4*(-/-) model but are thought to function through distinct mechanisms. Specifically, while the Hif-1α-independent benefits of chronic mild hypoxia are not well understood, the intervention is not thought to act via broad immune cell depletion [24]. Rather, hypoxia is generally thought to act by directly impacting mitochondrial function (ETC CI function, metabolic regulation at the mitochondria, etc). Accordingly, hypoxia is predicted to act upstream of immune activation in LS, preventing the induction of the immune-activating stress, the specific nature of which remains to be defined.

Here, without needing to resolve the proximal mechanisms of hypoxia, we provide evidence in support of this notion by demonstrating that cessation of hypoxia leads to a rapid induction and progression of disease in the *Ndufs4*(-/-), starkly contrasting with the slow development of disease in mice treated with an immune cell depleting drug. These findings have multiple mechanistic and therapeutic implications. In terms of disease mechanisms, they support the notion that disease onset is linked to development, with the pathway of immune activation developing postnatally and corresponding with disease onset (∼P37 in mice). As predicted by this model, mice removed from hypoxia after this age show a rapid induction (or de-repression) of immune activation leading to accelerated disease progression. In these animals, the immune initiating stress occurs in a system where the immune system and/or stress pathways are fully developed. In contrast, symptoms onset was delayed by 1-2 weeks post treatment cessation in pexidartinib treated animals, consistent with the known timing of innate immune cell repopulation following pharmacologic depletion [71, 72].

Regarding therapeutic development, these findings suggest that if hypoxia-related interventions are to be successful, caution must be taken to ensure that patients on hypoxia-related therapies do not experience lapses in treatment. Disease onset was extremely rapid in mice removed from hypoxia (within 1-2 days), suggesting even a single missed dose of a hypoxia-mimetic could lead to rapid immune induction and resultant pathology. Combination therapy with hypoxia and immune-targeting agents might provide a strategy which ensures patients receive the maximal benefits of hypoxia without the risk of rapid disease onset. Of course, elucidating the precise mechanisms of immune activation may lead to more robust and precise interventions.

Elucidating the specific mechanisms of innate immune activation should ultimately provide a full picture of the relationship between hypoxia and immune depletion, resolve key outstanding questions in the pathogenesis of LS, and provide novel targets for intervention.

## Methods

### Animals

Mice were kept at the SCRI on a standard 12-hour light-dark cycle with food and water provided *ad libitum*. Mice were sacrificed using cervical dislocation, blood collected, and brain was dissected into following regions: cerebellum, the rest of the cerebrum, and brainstem, and were snap frozen in liquid nitrogen and stored at -80°C.

### Longitudinal assessments of disease

Clasping was assessed by visual scoring, as previously described [31, 32]. As disease progresses in *Ndufs4*(KO)’s animals display intermittent/transient improvement of symptoms. Here, we report whether animals *ever* presented the symptom for two or more consecutive days, a criterion which minimizes spurious reporting. For observational assessments, lab-wide quality control discussions occurred frequently to ensure consistency between technicians/researchers, and staff contributed equally to each treatment group to minimize any potential bias between individuals. Reported day of cachexia onset in the *Ndufs4*(-/-) is the day of life when an individual animal’s weight peaks and progressive weight loss begins.

#### Pharmacologic interventions

Mouse chow was ground to powder and mixed with drugs (see table in Figure 1). 300mL of 1% agar melted in sterile water was added per kilogram powdered chow and the mixture was pelleted and incubated at 37°C for 3-5 hours until dry. Pellets were stored at 4°C short term (up to 30-days) or -20°C long-term (up to 6-months). We previously demonstrated that this processing has no impact on animal health, survival, or disease (4). Food consumption calculated based on data in Figure S2 – the value of (0.15g-food/g-mouse/day), the lower end of daily food consumption estimate, was used to ensure mice received adequate drug.

### Replicate numbers, controls, and drug administration

Animal numbers for each dataset are in figure legends. Whenever possible all datapoints are shown.

Assignment to control treatment groups was spread throughout the duration of these studies to ensure that no shifts in colony survival, behavior, etc, occurred during the course of these experiments. Accordingly, control treatment groups generally contain larger cohorts than the individual treatments.

Rapamycin was provided as ABI-009 (nab-rapamycin), an albumin encapsulated water-soluble formulation. ABI-009 was provided by Aadi LLC (17383 Sunset Blvd, Pacific Palisades, CA 90272) in lyophilized form. ABI-009 was resuspended to 1.2mg/mL rapamycin in 1XPBS. This solution was sterile-filtered and stored in aliquots at -80°C. ABI-009 was administered at 66µL/10g for a dose of 8mg/kg/day rapamycin, as in prior studies (4, 5).

### RNA extraction and reverse transcription

Brain tissue samples were homogenized using Dounce homogenizers. Briefly, each frozen brainstem tissue sample was dropped into 1 ml QIAzol solution before it was allowed to thaw, and homogenized by hand, then the solution was transferred to an Eppendorf tube and placed on dry ice. Samples were stored at -80°C until RNA extraction using Qiagen RNeasy Lipid Tissue Mini kit following manufacturer’s guidelines. Briefly, 200 µl chloroform was added to each homogenized sample, shaken vigorously and then centrifuged at 12,000 rcf for 15 minutes at 4°C. Aqueous phase containing RNA was transferred to a fresh tube with an equal volume of 70% ethanol added. Up to 700 µl was then added to the spin column and centrifuged at 8,000 rcf for 15 seconds at room temperature, with eluent discarded from the collection tube and repeated with the remainder of the solution. The column was then washed in a similar way with RW1 and RPE buffers twice, second time centrifuged for two minutes. Then the column was placed into a new collection tube and spun down again for an extra minute to ensure no traces of ethanol were transferred from the RPE buffer. RNase-free water was added to the column, which was placed into a new collection tube, and centrifuged for one minute to elute the RNA. RNA was quantified using NanoDrop.

1500 ng of total RNA was used as a template for reverse transcription utilizing SuperScript VILO cDNA Synthesis Kit (Invitrogen; catalog #11754-050) following manufacturer’s instructions. The cDNA product concentration from the reaction was determined using NanoDrop and all the samples were diluted to the same concentration for subsequent qRT-PCR.

### Quantitative real-time PCR

qRT-PCR was performed using Universal TaqMan MasterMix and TaqMan FAM probes against the genes of interest. All reactions were run together with a TaqMan VIC probe against *ActB* housekeeping gene to which all the data were normalized. A standard curve was included in every run. The following TaqMan probes were used: Mm03047343_m1 Cd68 FAM-MGB (250 rxns), Mm00488332_m1 Siglec1 FAM-MGB (250 rxns), Mm00550376_m1 Vwf FAM-MGB (250 rxns), Mm01242576_m1 Pecam1 FAM-MGB (250 rxns), Mm02619580_g1 Actb VIC-MGB (2900 rxns).

### NanoString analysis

RNA extracted from brainstem samples for subsequent NanoString was diluted in sterile water to yield the final concentration of 20 ng/µl and subjected to NanoString nCounter system using PanCancer Immune Profiling gene panel consisting of 850 genes, including housekeeping genes. The analysis pipeline was as follows: data (transcript counts per gene per sample) were normalized to the housekeeping gene that showed the least variability across all samples, *Eef1g*. Outliers were removed using the Grubb’s test (which can only identify up to one outlier per group) with alpha set to 0.05 using GraphPad Prism version 11.0.2. Discovery analysis was performed using multiple unpaired t-tests assuming normal distribution, same standard deviation between populations, and using a false discovery rate (FDR) approach. Specifically, the Benjamini-Hochberg test was used with FDR q=0.01.

## Supplemental Material

**Figure S1.**
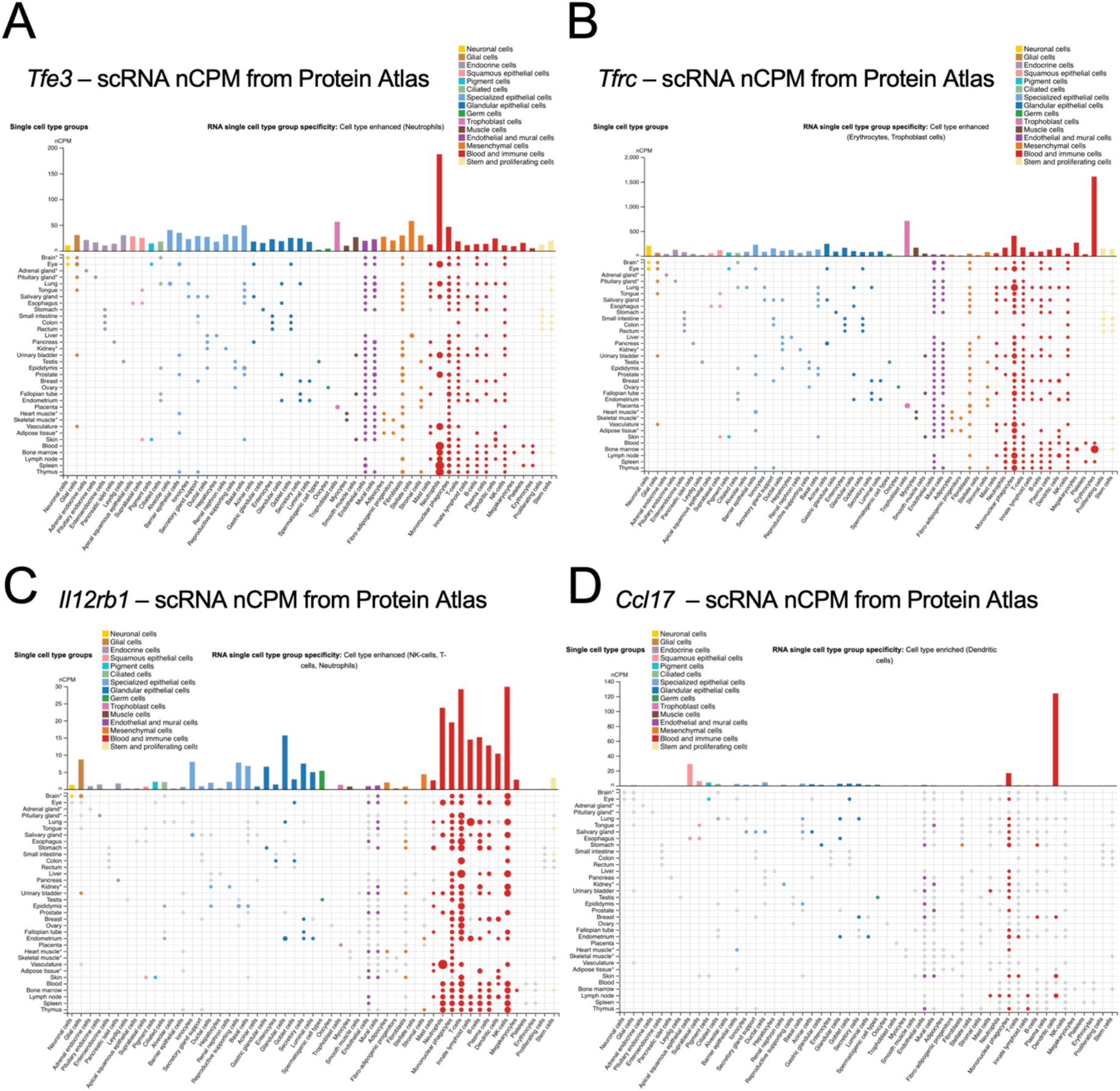
Human cell type specific expression of key factors upregulated in *Ndufs4*(-/-) brainstem at P23. Single cell RNA sequencing normalized counts per million (scRNA nCPM) data from Protein Atlas - see (Uhlen M et al., A genome-wide transcriptomic analysis of protein-coding genes in human blood cells. Science. (2019)) and (Karlsson M et al., A single-cell type transcriptomics map of human tissues. Sci Adv. (2021)), also cited in line in main text. Collected from proteinatlas.org on Aug 13, 2026. (A) *Tfe3*, (B) *Tfrc*, (C) *Il12rb1*, and (D) *Ccl17*. Cell and tissue types as indicated by text labels (axis) and color coding (inset keys).

**Figure S2.**
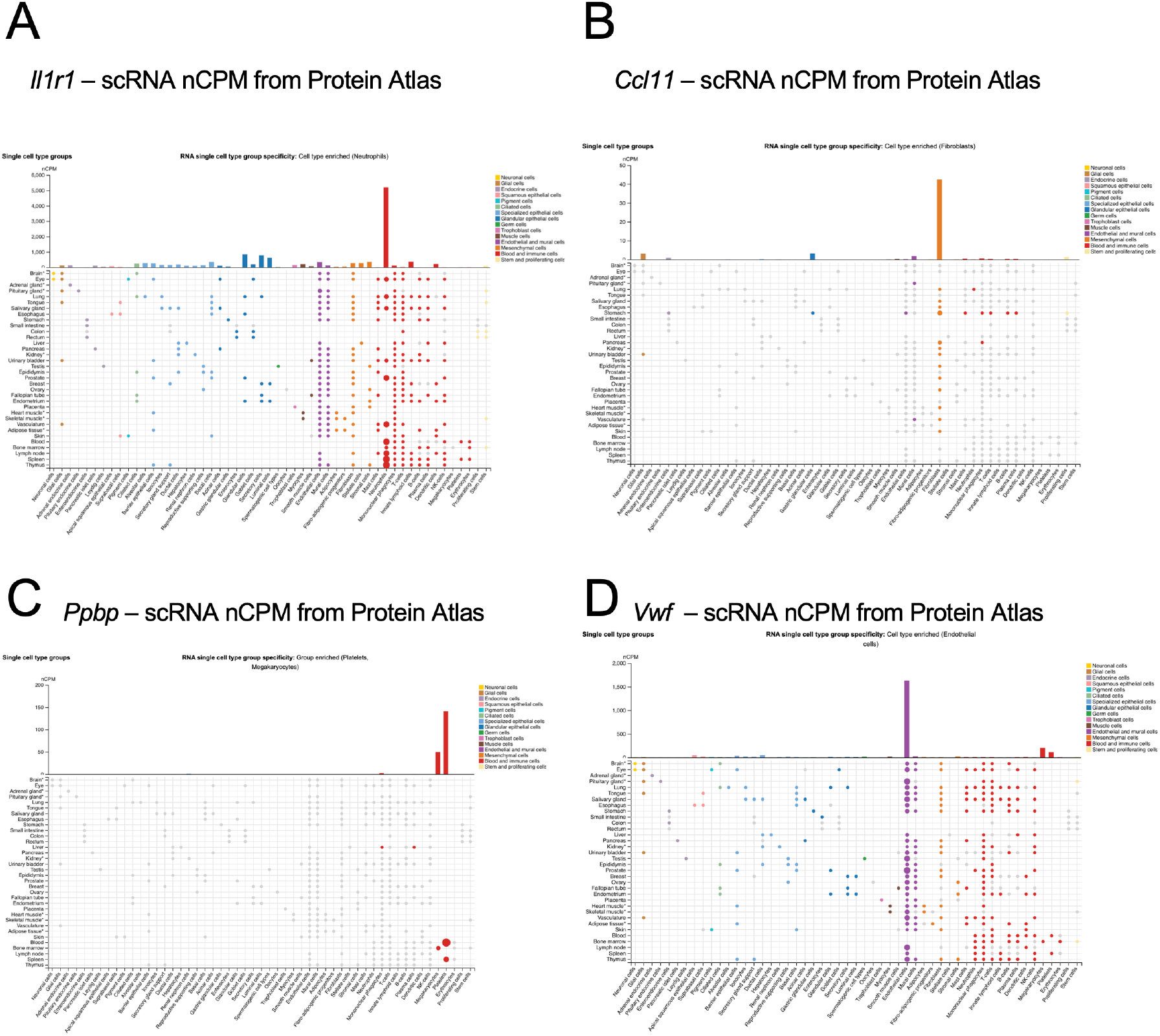
Human cell type specific expression of key factors increased in pexidartinib treated *Ndufs4*(-/-) brainstem at P45. Single cell RNA sequencing normalized counts per million (scRNA nCPM) data from Protein Atlas - see (Uhlen M et al., A genome-wide transcriptomic analysis of protein-coding genes in human blood cells. Science. (2019)) and (Karlsson M et al., A single-cell type transcriptomics map of human tissues. Sci Adv. (2021)), also cited in line in main text. (A) *Il1r1*, (B) *Ccl11*, (C) *Ppbp*, and (D) *Vwf*. Cell and tissue types as indicated by text labels (axis) and color coding (inset keys).

**Supplemental file: Mouse PanCancer Immune Panel** – NanoString panel details including target transcripts, sequences of probes, and list of targets associated with individual immune cell types. Information adapted from NanoString Technologies Whitepaper “Multiplexed Cancer Immune Response Analysis”, 2019, and nCounter® Mouse PanCancer Immune Profiling Panel - Gene and Probe Details.

**Supplemental table: Complete blood counts (CBC) from hypoxia cessation animals**. Units as shown in left column. WBC – white blood cells, LYM – lymphocytes, MON – monocytes, NEU – neutrophils, RBC – red blood cells, HGB – hemoglobin, HCT – hematocrit, MCV – mean cell volume, MCH - mean corpuscular hemoglobin, MCHC - mean corpuscular hemoglobin concentration, RDW - red cell distribution width, PLT – platelet, MPV – mean platelet volume, PCT – plateletcrit, PDW – platelet distribution width.

## Citations

1. Magro, G., V. Laterza, and F. Tosto, Leigh Syndrome: A Comprehensive Review of the Disease and Present and Future Treatments. Biomedicines, 2025. 13(3).

2. Alemao, N.N.G., et al., Leigh’s disease, a fatal finding in the common world: A case report. Radiol Case Rep, 2022. 17(9): p. 3321–3325.

3. Kruse, S.E., et al., Mice with mitochondrial complex I deficiency develop a fatal encephalomyopathy. Cell Metab, 2008. 7(4): p. 312–20.

4. Vafaee-Shahi, M., et al., Bilateral horizontal gaze palsy in an 8-year-old girl: A rare case with NDUFS4 gene mutation. Clin Case Rep, 2021. 9(9): p. e04748.

5. Petruzzella, V., et al., A nonsense mutation in the NDUFS4 gene encoding the 18 kDa (AQDQ) subunit of complex I abolishes assembly and activity of the complex in a patient with Leigh-like syndrome. Hum Mol Genet, 2001. 10(5): p. 529–35.

6. Ortigoza-Escobar, J.D., et al., Ndufs4 related Leigh syndrome: A case report and review of the literature. Mitochondrion, 2016. 28: p. 73–8.

7. Misceo, D., et al., Biallelic NDUFA4 Deletion Causes Mitochondrial Complex IV Deficiency in a Patient with Leigh Syndrome. Genes (Basel), 2024. 15(4).

8. Martin, M.A., et al., Leigh syndrome associated with mitochondrial complex I deficiency due to a novel mutation in the NDUFS1 gene. Arch Neurol, 2005. 62(4): p. 659–61.

9. Leshinsky-Silver, E., et al., NDUFS4 mutations cause Leigh syndrome with predominant brainstem involvement. Mol Genet Metab, 2009. 97(3): p. 185–9.

10. Budde, S.M., et al., Combined enzymatic complex I and III deficiency associated with mutations in the nuclear encoded NDUFS4 gene. Biochem Biophys Res Commun, 2000. 275(1): p. 63–8.

11. Anderson, S.L., et al., A novel mutation in NDUFS4 causes Leigh syndrome in an Ashkenazi Jewish family. J Inherit Metab Dis, 2008. 31 Suppl 2: p. S461–7.

12. Quintana, A., et al., Complex I deficiency due to loss of Ndufs4 in the brain results in progressive encephalopathy resembling Leigh syndrome. Proc Natl Acad Sci U S A, 2010. 107(24): p. 10996–1001.

13. Stokes, J.C., et al., Leukocytes mediate disease pathogenesis in the Ndufs4(KO) mouse model of Leigh syndrome. JCI Insight, 2022. 7(5).

14. Hanaford, A.R., et al., Peripheral macrophages drive CNS disease in the Ndufs4(-/-) model of Leigh syndrome. Brain Pathol, 2023: p. e13192.

15. Hanaford, A.R., et al., Disruption of adaptive immunity does not attenuate disease in the Ndufs4(-/-) model of Leigh syndrome. PLoS One, 2025. 20(6): p. e0324268.

16. Spencer, K.A., et al., Volatile anaesthetic toxicity in the genetic mitochondrial disease Leigh syndrome. Br J Anaesth, 2023. 131(5): p. 832–846.

17. Quintana, A., et al., Fatal breathing dysfunction in a mouse model of Leigh syndrome. J Clin Invest, 2012. 122(7): p. 2359–68.

18. Bolea, I., et al., Defined neuronal populations drive fatal phenotype in a mouse model of Leigh syndrome. Elife, 2019. 8.

19. Johnson, S.C., et al., Regional metabolic signatures in the Ndufs4(KO) mouse brain implicate defective glutamate/alpha-ketoglutarate metabolism in mitochondrial disease. Mol Genet Metab, 2020. 130(2): p. 118–132.

20. Hanaford, A.R., et al., Interferon-gamma contributes to disease progression in the Ndufs4(-/-) model of Leigh syndrome. Neuropathol Appl Neurobiol, 2024. 50(3): p. e12977.

21. Martin-Perez, M., et al., PKC downregulation upon rapamycin treatment attenuates mitochondrial disease. Nat Metab, 2020. 2(12): p. 1472–1481.

22. Ferrari, M., et al., Hypoxia treatment reverses neurodegenerative disease in a mouse model of Leigh syndrome. Proc Natl Acad Sci U S A, 2017. 114(21): p. E4241–E4250.

23. Jain, I.H., et al., Hypoxia as a therapy for mitochondrial disease. Science, 2016. 352(6281): p. 54–61.

24. Jain, I.H., et al., Leigh Syndrome Mouse Model Can Be Rescued by Interventions that Normalize Brain Hyperoxia, but Not HIF Activation. Cell Metab, 2019. 30(4): p. 824–832 e3.

25. Blume, S.Y., et al., HypoxyStat, a small-molecule form of hypoxia therapy that increases oxygen-hemoglobin affinity. Cell, 2025. 188(6): p. 1580–1588 e11.

26. Li, X., et al., Emerging roles of TFE3 in metabolic regulation. Cell Death Discov, 2023. 9(1): p. 93.

27. Karlsson, M., et al., A single-cell type transcriptomics map of human tissues. Sci Adv, 2021. 7(31).

28. Uhlen, M., et al., A genome-wide transcriptomic analysis of protein-coding genes in human blood cells. Science, 2019. 366(6472).

29. Sato, T., et al., Interferon regulatory factor-2 protects quiescent hematopoietic stem cells from type I interferon-dependent exhaustion. Nat Med, 2009. 15(6): p. 696–700.

30. Szklarczyk, D., et al., The STRING database in 2023: protein-protein association networks and functional enrichment analyses for any sequenced genome of interest. Nucleic Acids Res, 2023. 51(D1): p. D638–D646.

31. Johnson, S.C., et al., Dose-dependent effects of mTOR inhibition on weight and mitochondrial disease in mice. Front Genet, 2015. 6: p. 247.

32. Johnson, S.C., et al., mTOR inhibition alleviates mitochondrial disease in a mouse model of Leigh syndrome. Science, 2013. 342(6165): p. 1524–8.

33. Bornstein, R., et al., Differential effects of mTOR inhibition and dietary ketosis in a mouse model of subacute necrotizing encephalomyelopathy. Neurobiol Dis, 2022. 163: p. 105594.

34. Doyle, C.K., et al., Hyperconservation of the N-formyl peptide binding site of M3: evidence that M3 is an old eutherian molecule with conserved recognition of a pathogen-associated molecular pattern. J Immunol, 2003. 171(2): p. 836–44.

35. Hong, C.M., et al., Clinical Characteristics of Early-Onset and Late-Onset Leigh Syndrome. Front Neurol, 2020. 11: p. 267.

36. Antonicka, H., et al., Bi-allelic mutations in FASTKD5 are associated with cytochrome c oxidase deficiency and early-to late-onset Leigh syndrome. Am J Hum Genet, 2025. 112(7): p. 1699–1710.

37. Carli, S., et al., Natural History of Patients With Mitochondrial ATPase Deficiency Due to Pathogenic Variants of MT-ATP6 and MT-ATP8. Neurology, 2025. 104(7): p. e213462.

38. Liao, Y., et al., Adult-onset Leigh syndrome with recurrent seizures and peripheral neuropathy due to the S17CT > C mutation: a case report and literature review. BMC Neurol, 2025. 25(1): p. 128.

39. Hanaford, A. and S.C. Johnson, The immune system as a driver of mitochondrial disease pathogenesis: a review of evidence. Orphanet J Rare Dis, 2022. 17(1): p. 335.

40. Wang, W.H., et al., Studying Hair Growth Cycle and its Effects on Mouse Skin. J Invest Dermatol, 2023. 143(9): p. 1638–1645.

41. Santos-Lima, B., et al., The role of neutrophils in the dysfunction of central nervous system barriers. Front Aging Neurosci, 2022. 14: p. 965169.

42. Chakraborty, S., et al., A Brief Overview of Neutrophils in Neurological Diseases. Biomolecules, 2023. 13(5).

43. Dorward, D.A., et al., The role of formylated peptides and formyl peptide receptor 1 in governing neutrophil function during acute inflammation. Am J Pathol, 2015. 185(5): p. 1172–84.

44. Leslie, J., et al., FPR-1 is an important regulator of neutrophil recruitment and a tissue-specific driver of pulmonary fibrosis. JCI Insight, 2020. 5(4).

45. Kwon, W.Y., et al., Removal of circulating mitochondrial N-formyl peptides via immobilized antibody therapy restores sepsis-induced neutrophil dysfunction. J Leukoc Biol, 2024. 116(5): p. 1169–1183.

46. Johnson, S.C., et al., mTOR inhibitors may benefit kidney transplant recipients with mitochondrial diseases. Kidney Int, 2019. 95(2): p. 455–466.

47. Saemann, M.D., et al., The multifunctional role of mTOR in innate immunity: implications for transplant immunity. Am J Transplant, 2009. 9(12): p. 2655–61.

48. Patel, C.H. and J.D. Powell, More TOR: The expanding role of mTOR in regulating immune responses. Immunity, 2025. 58(7): p. 1629–1645.

49. Jones, G.E., et al., Requirement for PI 3-kinase gamma in macrophage migration to MCP-1 and CSF-1. Exp Cell Res, 2003. 290(1): p. 120–31.

50. Solinas, G. and B. Becattini, The role of PI3Kgamma in metabolism and macrophage activation. Oncotarget, 2017. 8(63): p. 106145–106146.

51. Hong, D.S., et al., Eganelisib, a First-in-Class PI3Kgamma Inhibitor, in Patients with Advanced Solid Tumors: Results of the Phase 1/1b MARIO-1 Trial. Clin Cancer Res, 2023. 29(12): p. 2210–2219.

52. Liu, M., et al., G protein-coupled receptor FPR1 as a pharmacologic target in inffammation and human glioblastoma. Int Immunopharmacol, 2012. 14(3): p. 283–8.

53. Vadas, O., et al., Molecular determinants of PI3Kgamma-mediated activation downstream of G-protein-coupled receptors (GPCRs). Proc Natl Acad Sci U S A, 2013. 110(47): p. 18862–7.

54. Piedra-Quintero, Z.L., et al., CD38: An Immunomodulatory Molecule in Inffammation and Autoimmunity. Front Immunol, 2020. 11: p. 597959.

55. Chini, E.N., CD38 as a regulator of cellular NAD: a novel potential pharmacological target for metabolic conditions. Curr Pharm Des, 2009. 15(1): p. 57–63.

56. Hogan, K.A., C.C.S. Chini, and E.N. Chini, The Multi-faceted Ecto-enzyme CD38: Roles in Immunomodulation, Cancer, Aging, and Metabolic Diseases. Front Immunol, 2019. 10: p. 1187.

57. Chini, C.C.S., et al., CD38 ecto-enzyme in immune cells is induced during aging and regulates NAD(+) and NMN levels. Nat Metab, 2020. 2(11): p. 1284–1304.

58. Schultz, M.B. and D.A. Sinclair, Why NAD(+) Declines during Aging: It’s Destroyed. Cell Metab, 2016. 23(6): p. 965–966.

59. Inoue, T., et al., Bone marrow stromal cell antigen-1 deficiency protects from acute kidney injury. Am J Physiol Renal Physiol, 2024. 326(2): p. F167–F177.

60. Lee, C.F., et al., Targeting NAD(+) Metabolism as Interventions for Mitochondrial Disease. Sci Rep, 2019. 9(1): p. 3073.

61. Chen, L., et al., Expression of the mouse MHC class Ib H2-T11 gene product, a paralog of H2-T23 (Qa-1) with shared peptide-binding specificity. J Immunol, 2014. 193(3): p. 1427–39.

62. Winther, M., et al., Formylated MHC Class Ib Binding Peptides Activate Both Human and Mouse Neutrophils Primarily through Formyl Peptide Receptor 1. PLoS One, 2016. 11(12): p. e0167529.

63. Beismann-Driemeyer, S. and R. Tampe, Function of the antigen transport complex TAP in cellular immunity. Angew Chem Int Ed Engl, 2004. 43(31): p. 4014–31.

64. Hermel, E., E. Grigorenko, and K.F. Lindahl, Expression of medial class I histocompatibility antigens on RMA-S mutant cells. Int Immunol, 1991. 3(4): p. 407–12.

65. Li, Y., et al., Targeting formyl peptide receptor 1 reduces brain inffammation and neurodegeneration. Science, 2025. 390(6774): p. eadq1177.

66. Zhangsun, Z., et al., FPR1: A critical gatekeeper of the heart and brain. Pharmacol Res, 2024. 202: p. 107125.

67. Schafer, D.P., et al., Microglia sculpt postnatal neural circuits in an activity and complement-dependent manner. Neuron, 2012. 74(4): p. 691–705.

68. Ding, X., et al., Loss of microglial SIRPalpha promotes synaptic pruning in preclinical models of neurodegeneration. Nat Commun, 2021. 12(1): p. 2030.

69. Ni, R.J., et al., Microglia-mediated inflammation and synaptic pruning contribute to sleep deprivation-induced mania in a sex-specific manner. Transl Psychiatry, 2025. 15(1): p. 285.

70. Geloso, M.C. and N. D’Ambrosi, Microglial Pruning: Relevance for Synaptic Dysfunction in Multiple Sclerosis and Related Experimental Models. Cells, 2021. 10(3).

71. Wang, Q., et al., Dynamic changes in microglia in the mouse hippocampus during administration and withdrawal of the CSF1R inhibitor PLX3397. J Anat, 2023. 243(3): p. 394–403.

72. van Rooijen, N., N. Kors, and G. Kraal, Macrophage subset repopulation in the spleen: differential kinetics after liposome-mediated elimination. J Leukoc Biol, 1989. 45(2): p. 97–104.

