## Supplementary figures and images for "Immune profiling and treatment cessation provide mechanistic insights into disease pathogenesis and considerations for translating pre-clinical strategies in Leigh syndrome"

### Figure S1

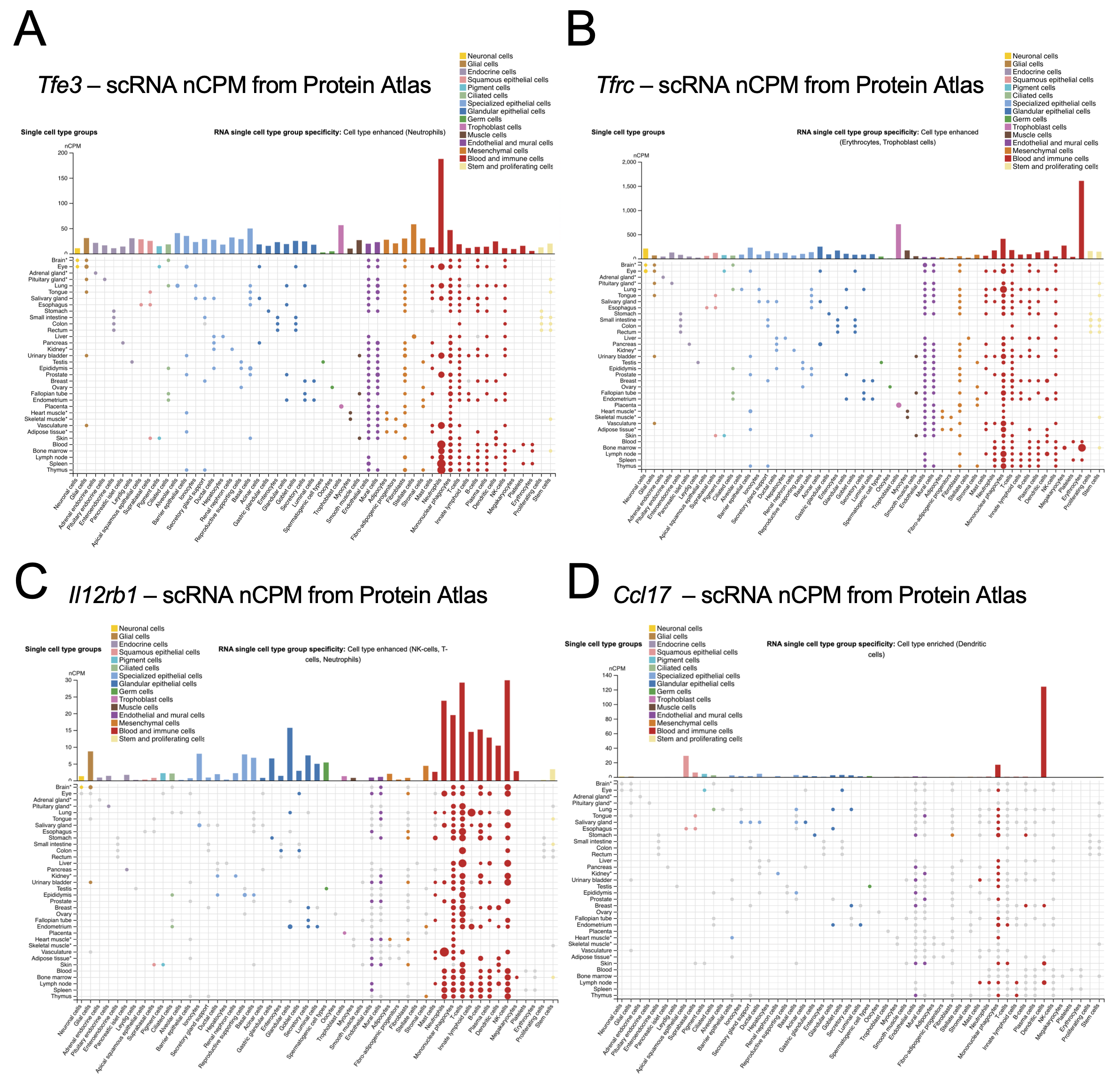

### Figure S2

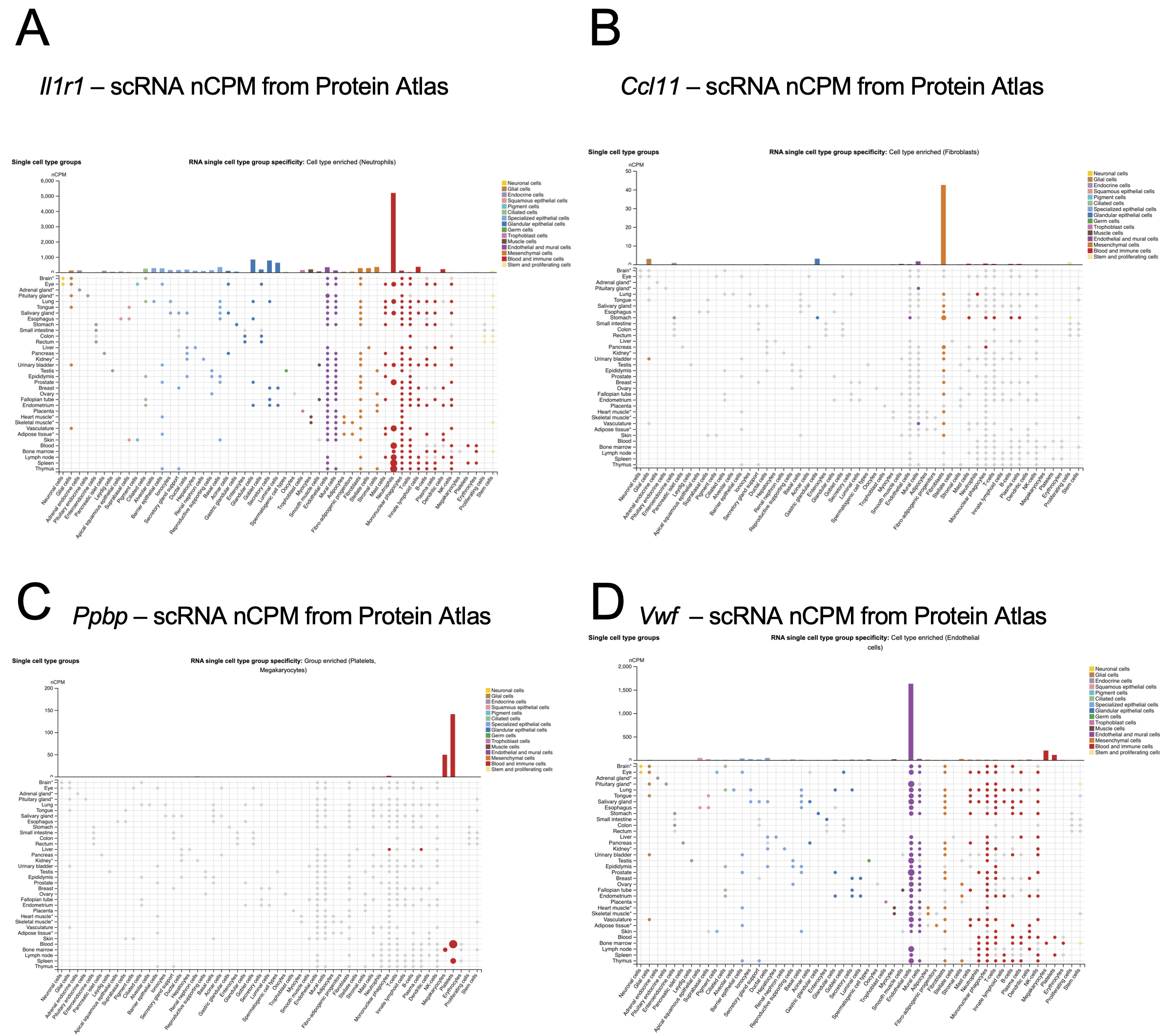
