## Supplemental table for "Immune profiling and treatment cessation provide mechanistic insights into disease pathogenesis and considerations for translating pre-clinical strategies in Leigh syndrome"

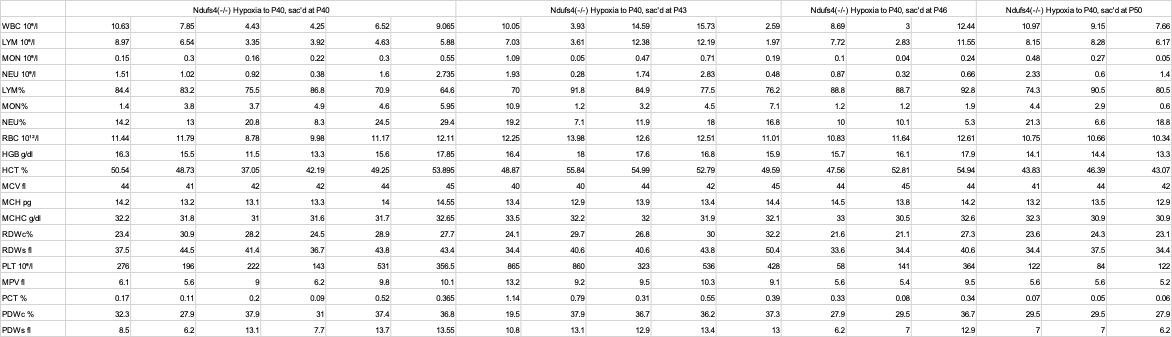

Table S1 – Complete blood counts (CBC) from hypoxia cessation animals. Units as shown in left column. WBC – white blood cells, LYM – lymphocytes, MON – monocytes, NEU – neutrophils, RBC – red blood cells, HGB – haemoglobin, HCT – haematocrit, MCV – mean cell volume, MCH - mean corpuscular hemoglobin, MCHC - mean corpuscular hemoglobin concentration, RDW - red cell distribution width, PLT – platelet, MPV – mean platelet volume, PCT – plateletcrit, PDW – platelet distribution width.
